# All-by-All Cytokine Receptor Pairing Network Unlocks Coding of Non-Natural T Cell States

**DOI:** 10.64898/2026.08.19.745857

**Authors:** Pingdong Tao, Kristin C.Y. Tsui, Yang Zhao, Hua Jiang, Zinaida Good, K. Christopher Garcia

**Affiliations:** Department of Molecular and Cellular Physiology, Stanford University School of Medicine, Stanford, CA, USA; Howard Hughes Medical Institute, Stanford University School of Medicine, Stanford, CA, USA; Department of Structural Biology, Stanford University School of Medicine, Stanford, CA, USA; Division of Immunology and Rheumatology, Stanford University School of Medicine, Stanford, USA; Division of Computational Medicine, Stanford University School of Medicine, Stanford, USA; Center for Cancer Cell Therapy, Stanford Cancer Institute, Stanford University, Stanford, USA

## Abstract

Cytokine receptor pairing rules, set by evolution, confine JAK-STAT signaling to a narrow region of a far larger combinatorial space. Of more than 1,200 pairings theoretically possible among the ∼36 JAK-associated human cytokine receptors, only ∼30–40 exist in nature. Using a double-orthogonal platform, we enforced pairings across the full all-by-all receptor matrix and resolved a fine-grained STAT atlas richer than the natural repertoire. Selected non-natural pairings generated emergent T cell states unpredictable from either parental receptor, with pairing orientation encoding signaling specificity. A synthetic IL-21R>IL-2Rβ pairing, but not its reciprocal, drove a cytotoxic Tc17-like state, whereas natural IL-9R/γc drove a Tc1 fate despite similar STAT activation, showing that rebalancing quantitative STAT combinatorics can modulate T cell fate. Recombining IL-31R, not expressed in T cells, with STAT-biased receptor partners generated diverse states, several with superior antitumor efficacy. These findings define a non-natural pairing code for engineering synthetic T cell fates.

## Introduction

T cells are remarkably plastic, reshaping their phenotypes in response to diverse environmental signals^1–5^. Cytokines are central instructors of this plasticity, regulating T cell differentiation and effector function by inducing cognate receptor dimerization and activating JAK-STAT signaling^6–10^. The functional logic of cytokine signaling has been framed around the dual properties of pleiotropy and redundancy, the abilities of a single cytokine to exert multiple distinct actions, and of different cytokines to elicit overlapping responses^11–14^. Decades of research has established that these properties arise in large part from the modular sharing of receptor subunits: common chains such as γc, gp130, βc, and IL-10Rβ are each used by multiple cytokines, while individual cytokines such as IL-2 and IL-4, can signal through more than one receptor complex, eliciting distinct outputs depending on which chains are engaged^15–17^. Viewed this way, receptor subunit sharing is a strategy that nature uses to expand functional diversity, generating new cytokine activities by recombining a limited set of receptor chains into alternative pairings, each wired to a distinct balance of downstream STATs.

This combinatorial reuse of receptor components, refined over evolution, implies that the natural repertoire of receptor pairings is only a small sampling of a larger space that the same chains could in principle form. Indeed, of the more than 1,200 receptor pairs theoretically possible among the ∼36 JAK-associated cytokine receptors encoded in the human genome, only ∼30-40 have been used in response to ∼50 endogenous cytokines over the course of evolution^18,19^, leaving the vast majority of the combinatorial signaling space unexplored. Efforts to diversify cytokine signaling inputs through affinity-tuned cytokine engineering^20–22^, synergistic cytokine combinations^23–26^, synthetic cytokine surrogate^27–29^, and orthogonal cytokine receptor engineering^30–33^ have begun to access small regions of this space, but the full landscape of possible receptor pairings, and the T cell states it could encode, has remained unmapped.

We reasoned that we could learn from the evolutionary examples of shared receptors and extend it by enforcing receptor pairings beyond those that naturally occur. The combinatorial logic underlying cytokine pleiotropy could be exploited to access STAT signaling programs and corresponding T cell functions that lie outside the natural repertoire. Because endogenous cytokine receptor expression on T cells is restricted, ectopic expression of orthogonal cytokine receptors offers a means to interrogate T cell plasticity across a broader range of signaling inputs. We previously found that selected orthogonal receptors, when enforced to pair with endogenous common γc, reprogrammed mouse T cells toward diverse phenotypes, from canonical Th2/Tc2 subsets induced by IL-4R to synthetic myeloid-like states induced by GCSFR^33^. More recently, an accompanying study using a double-orthogonal cytokine receptor platform mapped the signaling and transcriptional output of each individual cytokine receptor across the full natural repertoire, establishing a coarse-grained STAT map in which quantitative, combinatorial biases in STAT activation deterministically specify T cell fate^34^. That work showed that even subtle differences in STAT stoichiometry instruct profoundly distinct T cell states, but the natural receptor pairing repertoire defined by evolution places limits on functional diversity.

Here, we sampled the full combinatorial space of all cytokine receptor pairings to ask what lies beyond the natural landscape. Using the double-orthogonal architecture, we constructed a complete 36 × 36 matrix of chimeric cytokine receptor pairs, 1,296 combinations, the great majority of which are never formed by natural ligands. By fixing JAK-binding regions derived from IL-2Rβ and γc within the chimeric receptors to maximize the kinase activation reactions, we achieved programmable activation of arbitrary receptor pairs, yielding a fine-grained STAT signaling atlas. This all-by-all matrix revealed a fine-grained pattern of STAT activation whose combinatorial diversity exceeds that of the natural repertoire, including pairing-restricted STAT signatures unavailable to any natural ligand. Deeper characterization of select non-natural pairings revealed that minor differences in STAT ratios are sufficient to redirect T cells into unclassified functional states with distinct identities and antitumor properties, with the outcomes not predictable from either parental receptor alone. Collectively, these results establish that remixing cytokine receptor pairing unlocks a blueprint for fine-tuning JAK-STAT combinatorics to access T cell functional states beyond the reach of natural cytokine signaling.

## Results

### An all-by-all receptor pairing matrix reveals a fine-grained STAT signaling atlas

To sample the combinatorial signaling space by recombining receptor dimers, we adapted a previously developed double-orthogonal cytokine-receptor system based on the IL-2/IL-2Rβ/γc complex (companion manuscript^34^), which enables enforced signaling dimers between receptors that do not naturally pair **(Fig. 1A)**. By swapping the intracellular signaling domains of both IL-2Rβ and γc for those derived from arbitrary cytokine receptors, the double-orthogonal system can generate a densely interconnected ‘all-by-all’ synthetic receptor network of non-natural receptor pairings **(Fig. 1B, right)**, beyond the restricted receptor pairing landscape observed in nature **(Fig. 1B, left)**.

**Figure 1.**
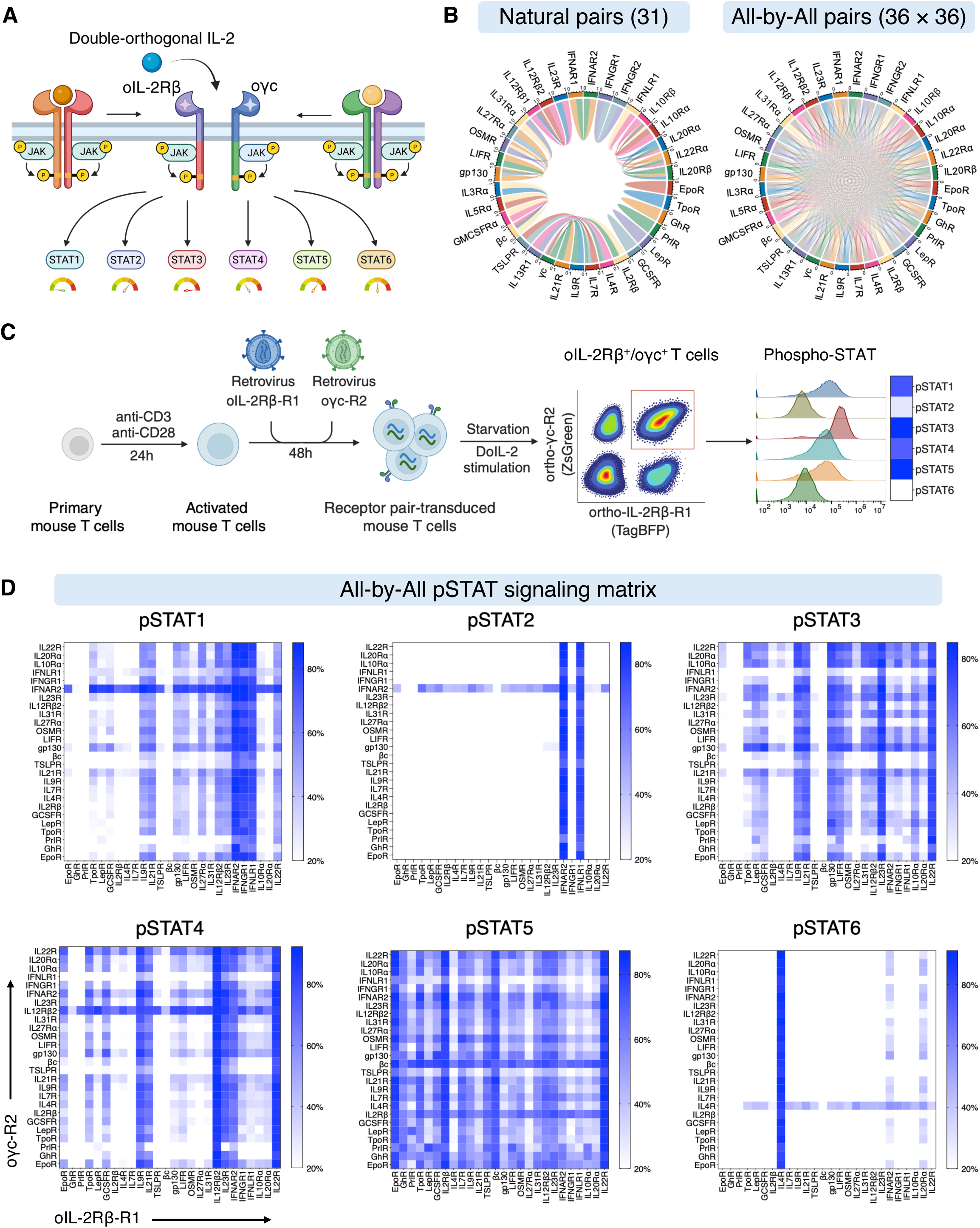
An all-by-all JAK-STAT signaling interactome across synthetic cytokine receptor pairs. (A) Schematic of the double-orthogonal receptor platform. The intracellular domain (ICD) of any cytokine receptor can be swapped into either orthogonal chain (oIL-2Rβ or oγc). Addition of double-orthogonal IL-2 (DoIL-2) forces the two chains together, activating downstream JAK-STAT signaling exclusively through the engineered pair. (B) Network diagrams comparing natural cytokine-mediated receptor pairings (∼31 pairs; left) with the all-by-all pairing network generated by the double-orthogonal system (36×36; right). (C) Experimental workflow for the all-by-all pSTAT signaling assay. Primary mouse T cells were sequentially transduced with oIL-2Rβ (TagBFP^+^) and oγc (ZsGreen^+^) chimeric receptors, starved, and stimulated with DoIL-2, then analyzed for phospho-STAT by flow cytometry in double-positive (TagBFP^+^/ZsGreen^+^) cells. (D) All-by-all pSTAT signaling heatmaps for STAT1-6. Each cell represents the pSTAT-positive frequency in double-positive T cells stimulated with DoIL-2, across all pairwise combinations of receptor ICDs carrying fixed IL-2Rβ- or γc-derived JAK-binding domains and natural STAT-recruiting regions. Rows represent oIL-2Rβ-fused ICDs (R1); columns represent oγc-fused ICDs (R2). See also **Figure S1.**

We first generated orthogonal chimeras containing the intracellular signaling domains (ICDs) of all 36 cytokine receptors in their native form. However, many non-native pairings elicited weak or undetectable STAT signaling, indicating that enforced dimerization alone is insufficient to drive productive signaling across arbitrary combinations **(Fig. 1C, Fig. S1A-B)**. We reasoned that this failure reflected inefficient JAK transphosphorylation and activation by some non-natural receptor pairings. To overcome this and achieve robust signaling from all possible receptor pairings, we employed a JAK-binding domain (Box region) fixation strategy^35–37^. In the optimized orthogonal receptor system, the Box region of each receptor was substituted with that derived from IL-2Rβ and γc^38,39^, ensuring that orthogonal ligand-induced receptor dimerization reliably assembles a naturally active JAK1/JAK3 kinase pair (in this case for IL-2) and enables downstream STAT phosphorylation regardless of the ICDs being tested **(Fig. S1C)**. This design exploits the fact that the γc/JAK3 node has evolved for substrate promiscuity as the shared signaling hub of the γc family, making JAK1/JAK3 transphosphorylation highly permissive and ideally suited for the all-by-all matrix^40,41^.

Using this optimized system, we profiled the STAT signaling matrix across all pairings of the 26 receptors containing STAT-binding motifs (26 × 26), omitting 10 receptors lacking apparent STAT binding motifs, and generated binary phosphorylation heatmaps across six STATs **(Fig. 1C-D, Fig. S1D)**. Nearly every receptor formed a functional signaling dimer with receptors beyond its native pairing, albeit with varying strength of pSTAT activation. The resulting atlas recovered expected specificities as internal validation: STAT2 activation was largely restricted to IFNAR2 and IFNLR1, and STAT6 was driven mainly by IL-4R, while revealing broad, promiscuous STAT5 activation and receptor-specific, tightly co-activated STAT1/3/4 patterns **(Fig. 1D)**. The same ICD consistently produced stronger STAT phosphorylation in the oIL-2Rβ arm (site 1, which binds to IL-2 first) than the oγc arm (site 2, which binds to IL-2 second) of the receptor heterodimer^42^ **(Fig. S1E-F)**, revealing an intrinsic signaling asymmetry between the two chains that can tunes STAT activation ratios.

In sum, this expanded atlas of STAT activation profiles provides a blueprint from which non-natural pairings with defined STAT signatures can be selected for functional interrogation.

### Reconstructing IL-9R-like programs through non-natural IL-2Rβ–IL-21R heterodimers

In our first experiment, we asked whether a non-natural pairing could be used to recapitulate the signaling and function of a natural cytokine receptor using a different combination of receptor chains, that could additively reconstruct the native STAT signature^16^. IL-2Rβ drives STAT5 through its natural partner γc; IL-21R drives STAT1/3/4 through γc. Forcing these two chains together into an IL-2Rβ–IL-21R heterodimer, bypassing γc, could in principle generate a hybrid STAT output with features from each receptor. We tested both orientations where each receptor was either in the site 1 or the site 2 position (IL-2Rβ/IL-21R or IL-21R/IL-2Rβ, respectively) alongside the natural IL-2Rβ/γc and IL-21R/γc pairings, and included IL-9R/γc as a natural benchmark because our previous data showed that it activates a broadly similar STAT1/3/4/5 profile and enhances antitumor T cell activity **(Fig. 2A, Fig. S2A)**^32,43,44^. Both reciprocal synthetic pairings combined STAT1/3/4 signaling derived from IL-21R with STAT5, MAPK, and PI3K-Akt-mTOR signaling from IL-2Rβ **(Fig. 2B, Fig. S2B-C)**. IL-9R/γc shows a comparable STAT profile but engaged these non-STAT pathways only weakly, providing a first indication that the synthetic and natural routes to a shared STAT signature are not signaling-equivalent.

**Figure 2.**
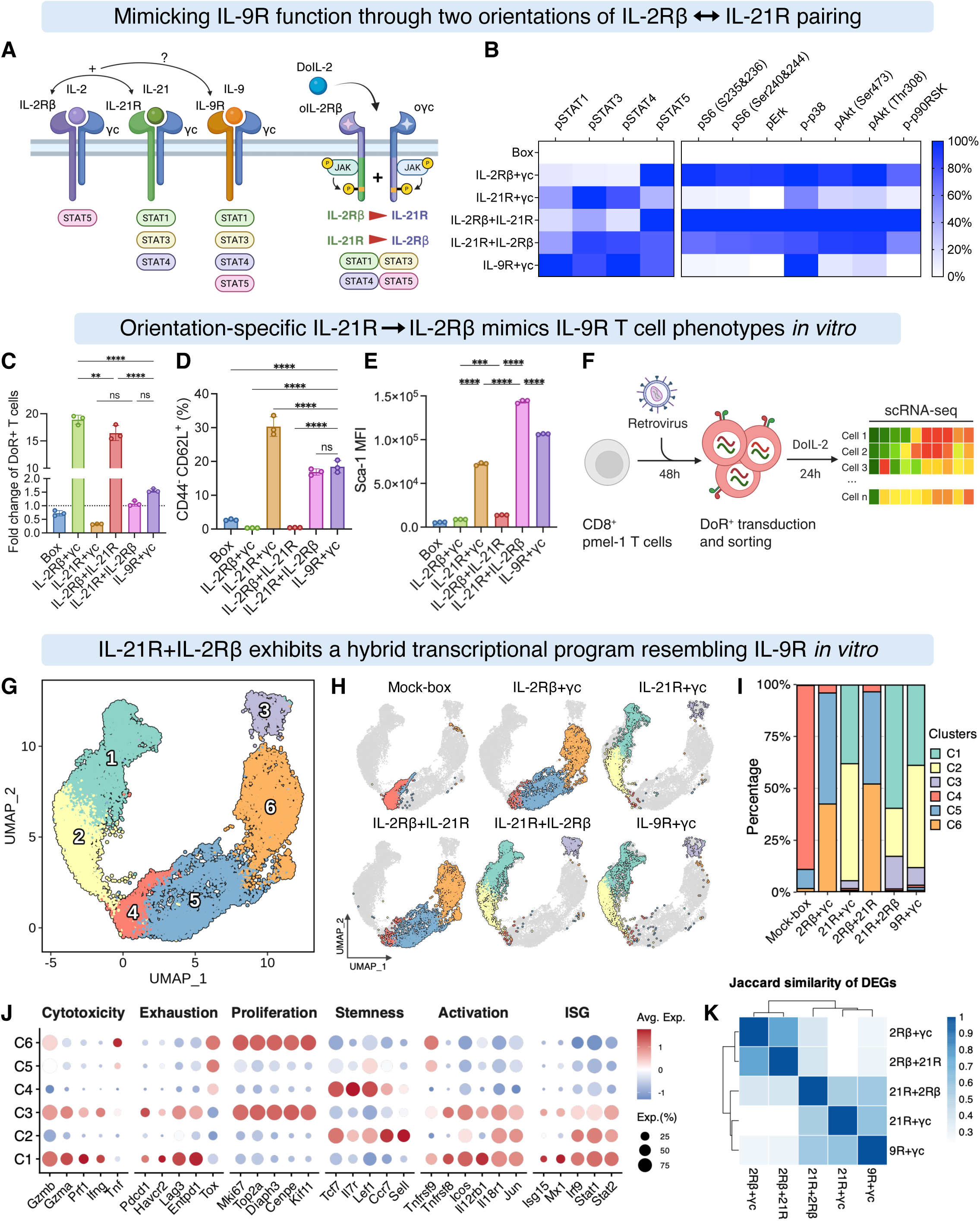
Non-natural IL-2Rβ–IL-21R pairings generate orientation-dependent hybrid signaling and emergent transcriptional programs. (A) Schematic of receptor pairings tested, including natural γc family heterodimers (IL-2Rβ/γc, IL-21R/γc, IL-9R/γc) and two reciprocal synthetic heterodimers (IL-2Rβ/IL-21R and IL-21R/IL-2Rβ), with their predicted downstream STAT pathways indicated. (B) Heatmap of normalized STAT and non-STAT phosphoprotein activities for each receptor pairing in Pmel-1 T cells stimulated with DoIL-2 (100 nM, 30 min). A truncated oIL-2Rβ/oγc construct retaining only the JAK-binding domain (Box control) was used as a signaling-null reference. Values represent normalized median fluorescence intensity (MFI). (C) *In vitro* proliferative expansion of Pmel-1 T cells expressing each receptor pairing. Cells were cultured with DoIL-2 (100 nM) for 3 days; orthogonal receptor-expressing T cells were counted by flow cytometry before and after culture. (D-E) Stem cell-like memory T (Tscm) cell markers after DoIL-2-stimulated expansion: CD44^−^CD62L^+^ frequency (D), Sca-1 MFI (E). (F) scRNA-seq workflow: sorted Pmel-1 CD8+ T cells transduced with each receptor pairing were stimulated with DoIL-2 (100 nM, 24 h) and processed for library generation using a plate-based combinatorial barcoding platform. (G) UMAP of 18,711 T cells from *in vitro* stimulation in (F). Six transcriptionally distinct clusters (C1-C6) are color-coded and annotated by dominant gene expression programs. (H) Per-condition UMAP distributions showing cluster occupancy for each receptor pairing. (I) Stacked bar plot showing the proportion of cells across annotated clusters. (J) Expression of key marker genes associated with distinct T cell functional states across annotated clusters. (K) Pairwise transcriptional similarity heatmap across all receptor pairings, calculated using the Jaccard similarity index of differentially expressed genes (log2FC > 0.5 vs. Box control). Data in (C-E) represent three biological replicates, mean ± SD. One-way ANOVA with Tukey’s post-test. ns, not significant; **p < 0.01; ***p < 0.001; ****p < 0.0001. See also **Figure S2.**

These signaling differences produced distinct T cell phenotypes *in vitro*. IL-2Rβ/γc drove robust proliferative expansion, whereas IL-21R/γc failed to expand CD8^+^ T cells at all (**Fig. 2C**). The two synthetic orientations diverged between these extremes: IL-2Rβ/IL-21R retained modest proliferation, whereas IL-21R/IL-2Rβ increased cell recovery without substantial proliferation, closely resembling IL-9R/γc. For stem-like memory T (Tscm) cells (CD44⁻CD62L⁺, Sca-1⁺), IL-21R/γc gave the highest frequency, but IL-21R/IL-2Rβ and IL-9R/γc generated the greatest absolute numbers and highest Sca-1 expression of this therapeutically important population **(Fig. 2D-E)**. Thus, the reciprocal pairings reconstructed distinct hybrid programs positioned between their parental receptors, with IL-21R/IL-2Rβ most closely, but not identically, approximating IL-9R/γc.

Single-cell RNA sequencing of 18,711 T cells across these pairing conditions (*in vitro* stimulation, 24h) (**Fig. 2F-I**) resolved six transcriptionally distinct clusters encompassing naïve, activated, proliferative, interferon-stimulated, exhaustion-associated, and cytotoxic programs (**Fig. 2J**). A naïve-like cluster (C4), marked by *Tcf7*, *Ccr7*, *Il7r*, and *Lef1*, was enriched in Box control (no ICD) cells. The remaining clusters split along two axes: IL-21R/γc, IL-21R/IL-2Rβ, and IL-9R/γc cells predominantly occupied clusters C1-C3, which exhibited more activated and cytotoxic phenotypes (**Fig. 2G**, **Fig. S2D-E**), while IL-2Rβ/γc and IL-2Rβ/IL-21R cells were enriched in the more proliferative clusters C5-C6. An intermediate hybrid cluster (C3), who’s differentially expressed genes overlapped >90% with either C1 or C6 (**Fig. S2F**), was selectively enriched in IL-21R/IL-2Rβ, and IL-9R/γc.

Although IL-2Rβ/IL-21R and IL-21R/IL-2Rβ contain identical receptor components, they elicited markedly distinct transcriptional programs, confirming the directional asymmetry to non-natural pairing. Genome-wide DEG similarity placed IL-2Rβ/IL-21R nearest the canonical IL-2Rβ/γc pairing, whereas IL-21R/IL-2Rβ aligned with IL-21R/γc and IL-9R/γc **(Fig. 2K, Fig. S2G)**. Consistently, GO^45^ biological-process enrichment showed that IL-21R/IL-2Rβ blends cell-cycle and RNA-metabolic programs characteristic of IL-2Rβ/γc with the T cell activation and differentiation programs of IL-21R/γc **(Fig. S2H)**.

Together, these data establish that receptor orientation, which manifests as different dimer geometries that skew the JAK/STAT substrate access, not composition alone, shapes signaling output and cell state, and that IL-21R/IL-2Rβ most closely recapitulates the hallmarks of IL-9R/γc through an entirely non-natural configuration. The significance of this result is that non-natural pairing signals can be approximated through remixing natively expressed receptors on T cells.

### Rewired IL-2Rβ–IL-21R heterodimers enhance adoptive T cell therapy

We next asked whether the signaling programs instigated by non-natural receptor pairings translate into improved antitumor activity *in vivo*. To minimize potential immunogenicity associated with human receptor extracellular domains, we rebuilt all receptor pairings using a fully murine orthogonal receptor system **(Fig. S3A)** and tested them in gp100-specific Pmel-1 CD8⁺ T cells against B16F10 melanoma, using MSA-fused mouse DoIL-2 for selective *in vivo* stimulation. We used two settings of increasing stringency: lymphodepleted mice, in which irradiation clears space for transferred cells^46,47^ **(Fig. 3A-C)**, and non-lymphodepleted mice, a more stringent test of T cell fitness **(Fig. 3D-F)**. All pairings were well tolerated **(Fig. S3B)**.

**Figure 3.**
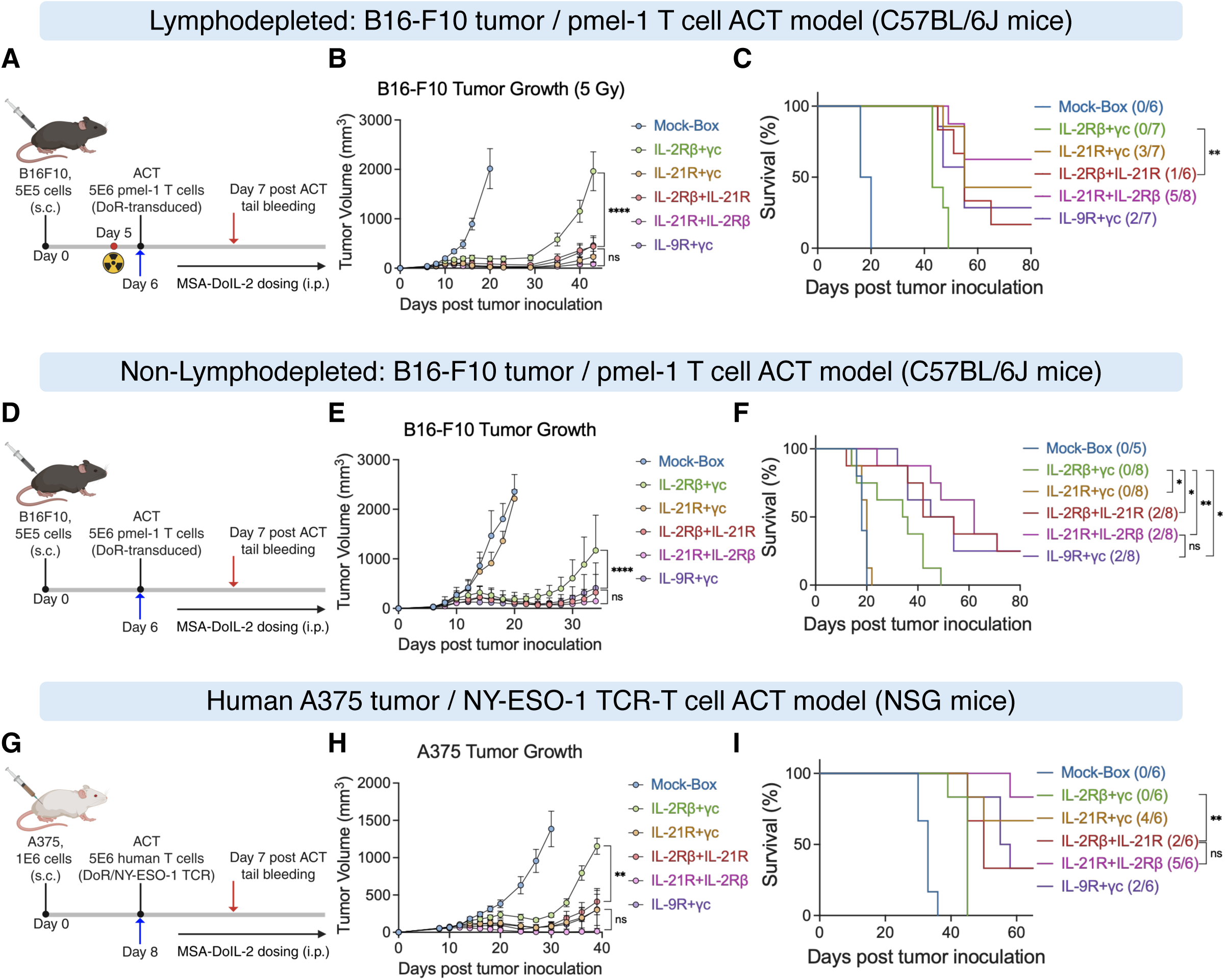
Non-natural IL-2Rβ–IL-21R heterodimers enhance adoptive T cell therapy in mouse and human tumor models. (A-C) Lymphodepleted syngeneic ACT model. Female C57BL/6 mice bearing subcutaneous B16F10 tumors were sublethally irradiated (5 Gy, day 5), then received retro-orbital transfer of 5×10⁶ Pmel-1 CD8^+^ T cells transduced with indicated receptor pairings (day 6), followed by intraperitoneal MSA-mouse DoIL-2 (15 μg, every other day through day 20; n = 6-8 per group). (A) Experimental timeline. (B) Average B16F10 tumor growth curves. (C) Survival curves. (D-F) Non-lymphodepleted syngeneic ACT model. As in (A-C) but without irradiation preconditioning (n = 5-8 per group). (D) Experimental timeline. (E) Average tumor growth curves. (F) Survival curves. (G-I) Human xenograft model. Male and female NSG mice bearing subcutaneous A375 tumors received retro-orbital transfer of 5×10⁶ human CD3^+^ T cells co-transduced with NY-ESO-1 TCR (1G4) and indicated receptor pairings (day 8), followed by MSA-human DoIL-2 (15 μg, every other day through day 22; n = 6 per group). (G) Experimental timeline. (H) Average A375 tumor growth curves. (I) Survival curves. Statistical tests: two-way ANOVA (B, E, H); log-rank Mantel-Cox (C, F, I). ns, not significant; *p < 0.05; **p < 0.01; ****p < 0.0001. See also **Figure S3.**

In the lymphodepleted mice, all receptor pairings delayed tumor growth versus Box control (**Fig. 3B**; individual tumor curves in **Fig. S3C**), but, surprisingly, efficacy and expansion were decoupled. IL-2Rβ/γc produced the highest peripheral T cell counts after transfer (**Fig. S3E**) yet achieved no complete responses (0/7 mice). By contrast, IL-21R/IL-2Rβ cured 5/8 mice, IL-21R/γc cured 3/7, IL-9R/γc cured 2/7, and IL-2Rβ/IL-21R cured 1/6 (**Fig. 3C**). Receptor pairings that mediated complete responses consistently exhibited higher Sca-1 expression on transferred T cells (**Fig. S3F**), indicating that therapeutic efficacy was more closely associated with acquisition of a stem-like phenotype than with the extent of peripheral expansion.

The functional differences among receptor pairings became even more pronounced in non-lymphodepleted mice. IL-21R/γc failed almost entirely: transferred cells were undetectable in blood (**Fig. S3G-H**) and mice showed no benefit beyond Box control (**Fig. 3E-F**). IL-2Rβ/γc delayed tumor growth but produced no cures. Strikingly, both orientations of the synthetic IL-2Rβ–IL-21R pairing, as well as IL-9R/γc, maintained curative efficacy comparable to that observed in lymphodepleted mice despite reduced peripheral T cell numbers (**Fig. 3E-F**; individual tumor curves in **Fig. S3D**), demonstrating that their superior therapeutic efficacy does not depend on lymphodepleting preconditioning.

These findings translated to a human xenograft model in which NY-ESO-1 TCR-T cells^48^ carrying human receptor orthologs were tested against A375 melanoma tumors in NSG mice (**Fig. 3G**). Human receptor pairings recapitulated the STAT profiles of their murine counterparts (**Fig. S3I**). Again, IL-2Rβ/γc was the only pairing failing to achieve complete responses, while both synthetic IL-2Rβ–IL-21R pairings and natural IL-21R/γc and IL-9R/γc all achieved cures (**Fig. 3H-I**; individual tumor curves in **Fig. S3J**), with IL-21R/IL-2Rβ giving the highest complete response rate among all conditions. CD62L expression on human transferred T cells (**Fig. S3K-L**) mirrored murine Sca-1 trends, confirming that stem-like programming is a conserved feature of the most therapeutically effective pairings.

These results show that non-natural, orientation-dependent IL-2Rβ–IL-21R pairings rebalance T cell expansion, persistence, and antitumor efficacy across multiple models. To ask whether these receptor programs drive T cells into convergent or distinct states within the tumor microenvironment, we next turned to single-cell profiling of tumor-infiltrating T cells to gain deeper insight into their cellular identities.

### IL-21R/IL-2Rβ and IL-9R/γc pairings drive distinct Tc17 and Tc1 programs *in vivo*

Although IL-21R/IL-2Rβ and IL-9R/γc exhibited similar *in vitro* STAT profiles and broadly comparable antitumor efficacy, they nevertheless showed subtle differences in peripheral T cell expansion and cellular phenotypes *in vivo*. To determine whether these differences reflected distinct intratumoral T cell states, we performed scRNA-seq on TILs from non-lymphodepleted B16F10 tumors after five DoIL-2 doses **(Fig. 4A)**. Nine transcriptional clusters were resolved across all of the conditions (**Fig. 4B, Fig. S4A**), with each pairing occupying a distinct cluster distribution (**Fig. 4C-D**). *In vivo* transcriptional relationships broadly mirrored *in vitro* patterns: IL-2Rβ/IL-21R TILs were closest to IL-2Rβ/γc, while IL-21R/IL-2Rβ TILs were closest to IL-21R/γc, though cluster distributions differed substantially between conditions. Marker gene expression further illustrated the distinct functional identities across receptor pairings, spanning memory, exhaustion, effector, Tc1, Tc17, and proliferative programs (**Fig. 4E**).

**Figure 4.**
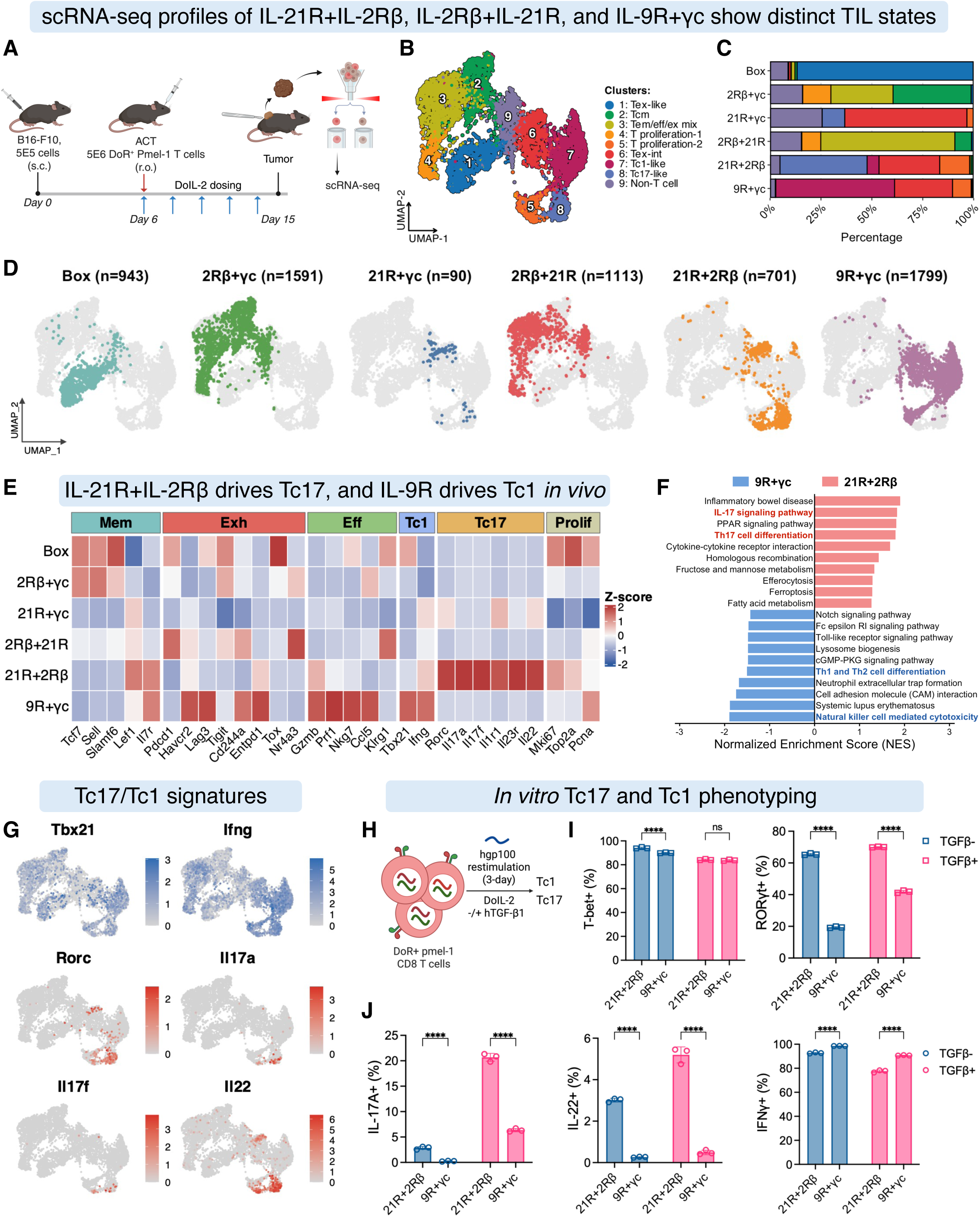
IL-21R/IL-2Rβ and IL-9R/γc drive distinct Tc17 and Tc1 tumor-infiltrating T cell states. (A) Experimental design for TIL scRNA-seq. Non-lymphodepleted C57BL/6 mice bearing B16F10 tumors received ACT of Pmel-1 T cells expressing indicated receptor pairings (day 6), followed by five doses of DoIL-2 (days 6-14). Thy1.1^+^CD8^+^OrthoR^+^ tumor-infiltrating T cells were sorted on day 15 for scRNA-seq using a plate-based combinatorial barcoding platform. (B) UMAP of sorted TILs across all six receptor pairing conditions. Nine transcriptionally distinct clusters are color-coded and annotated. (C) Stacked bar plots showing the proportion of cells per annotated cluster for each receptor pairing. (D) Per-condition UMAPs showing cluster distributions, with recovered T cell numbers indicated for each group. (E) Heatmap of selected marker genes grouped by functional category (memory, exhaustion, effector, Tc1, Tc17, proliferating) across all receptor pairings. Values are Z-scores. (F) KEGG pathway enrichment comparing IL-21R/IL-2Rβ and IL-9R/γc. Normalized enrichment scores (NES) are shown; positive values indicate enrichment in IL-21R/IL-2Rβ, negative in IL-9R/γc. (G) UMAP expression of Tc1-associated (*Tbx21*, *Ifng*) and Tc17-associated (*Rorc*, *Il17a*, *Il17f*, *Il22*) marker genes. Color intensity indicates normalized expression. (H) Schematic for *in vitro* Tc1/Tc17 validation. Pmel-1 T cells expressing IL-21R/IL-2Rβ or IL-9R/γc were restimulated with gp100 peptide (100 nM) and DoIL-2 (100 nM) ± TGF-β1 (10 nM) for 3 days, then analyzed for transcription factor expression and cytokine production. (I-J) Frequencies of T-bet+ and RORγt+ cells (I) and IL-17A+, IL-22+, and IFN-γ+ cells (J) by intracellular staining. Data represent three biological replicates, mean ± SD. Two-way ANOVA with Tukey’s post-test. ns, not significant; ****p < 0.0001. See also **Figure S4.**

Relative to Box control, IL-21R/IL-2Rβ and IL-9R/γc shared a broad activation program, characterized by coordinated upregulation of cytotoxicity, terminal differentiation, and activation genes, together with downregulation of stemness **(Fig. S4B-C)**. Directly comparing the two, however, revealed a profound divergence in lineage identity. IL-9R/γc selectively induced a canonical Tc1/NK-like cytotoxic program (*Gzma/c/f/g*, *Nkg7*, *Klrd1*, *Klre1*, *Tbx21*, high *Ifng*), whereas IL-21R/IL-2Rβ induced a Tc17 program (*Rorc*, *Il17a*, *Il17f*, *Il22*, *Il23r*, *Il1r1*, and multiple MAPK-family genes) **(Fig. S4D)**. KEGG^49^ pathway analysis supported these lineage identities: IL-9R/γc was enriched for NK cell-mediated cytotoxicity and Th1/Th2 differentiation, while IL-21R/IL-2Rβ was enriched for IL-17 signaling and Th17 differentiation (**Fig. 4F, Fig. S4E**). UMAP visualization of lineage markers further confirmed their segregation, with *Tbx21* and *Ifng* highest in IL-9R/γc clusters and *Rorc*, *Il17a*, *Il17f*, and *Il22* highest in IL-21R/IL-2Rβ clusters (**Fig. 4G**, **Fig. S4G**).

*In vitro* differentiation assays validated these transcriptional identities experimentally (**Fig. 4H**). After three days of restimulation with or without TGF-β, IL-21R/IL-2Rβ T cells showed markedly induced RORγt, while T-bet remained high in both conditions (>90% in each) (**Fig. 4I, Fig. S4H**). IL-21R/IL-2Rβ T cells robustly produced IL-17A and IL-22, further enhanced by TGF-β, while maintaining IFN-γ production at levels below but comparable to IL-9R/γc (**Fig. 4J, Fig. S4I**). IL-21R/IL-2Rβ therefore generates a hybrid Tc17 state that retains substantial cytotoxic function.

These results show that two receptor pairings with comparable, but non-identical, STAT activation profiles can drive T cells into fundamentally distinct functional identities *in vivo*, highlighting the sensitivity of T cell fate to STAT biases. The determining variable is quantitative: modestly higher STAT1 and STAT4 activation in IL-9R/γc versus IL-21R/IL-2Rβ (**Fig. S4F**) is sufficient to redirect fate from Tc17-like toward canonical Tc1. Quantitative STAT balance governs T cell fate choice, a principle we next exploited by engineering STAT combinations absent from any natural receptors.

### Rewiring a shared IL-31R scaffold with STAT-biased partners generates diverse antitumor programs

Having shown that non-natural pairings of T cell surface-expressed receptors can generate emergent states, we next asked whether we could go further and access STAT combinations absent from the endogenous receptor repertoire by building them onto a new common receptor scaffold. We chose IL-31R for this purpose because it is not naturally expressed on T cells^50,51^, and its natural pairing with OSMR drives predominantly STAT3/STAT5 while leaving STAT1, STAT2, STAT4, and STAT6 largely unengaged.

We paired IL-31R with five alternative receptor partners that it does not naturally engage with on cells, each contributing a distinct STAT axis: IFNAR2 (STAT1/2), IL-21R (STAT3), IL-12Rβ2 (STAT4), IL-2Rβ (STAT5), and IL-4R (STAT6) **(Fig. 5A)**. The STAT output of each partner receptor alone (i.e. paired with truncated IL-2Rβ or γc that has no STAT binding sites) **(Fig. 5B, Fig. S5A)** and of each rewired heterodimer pairings **(Fig. 5C, Fig. S5B),** confirmed that every combination produced a new STAT signature distinct from any natural receptor.

**Figure 5.**
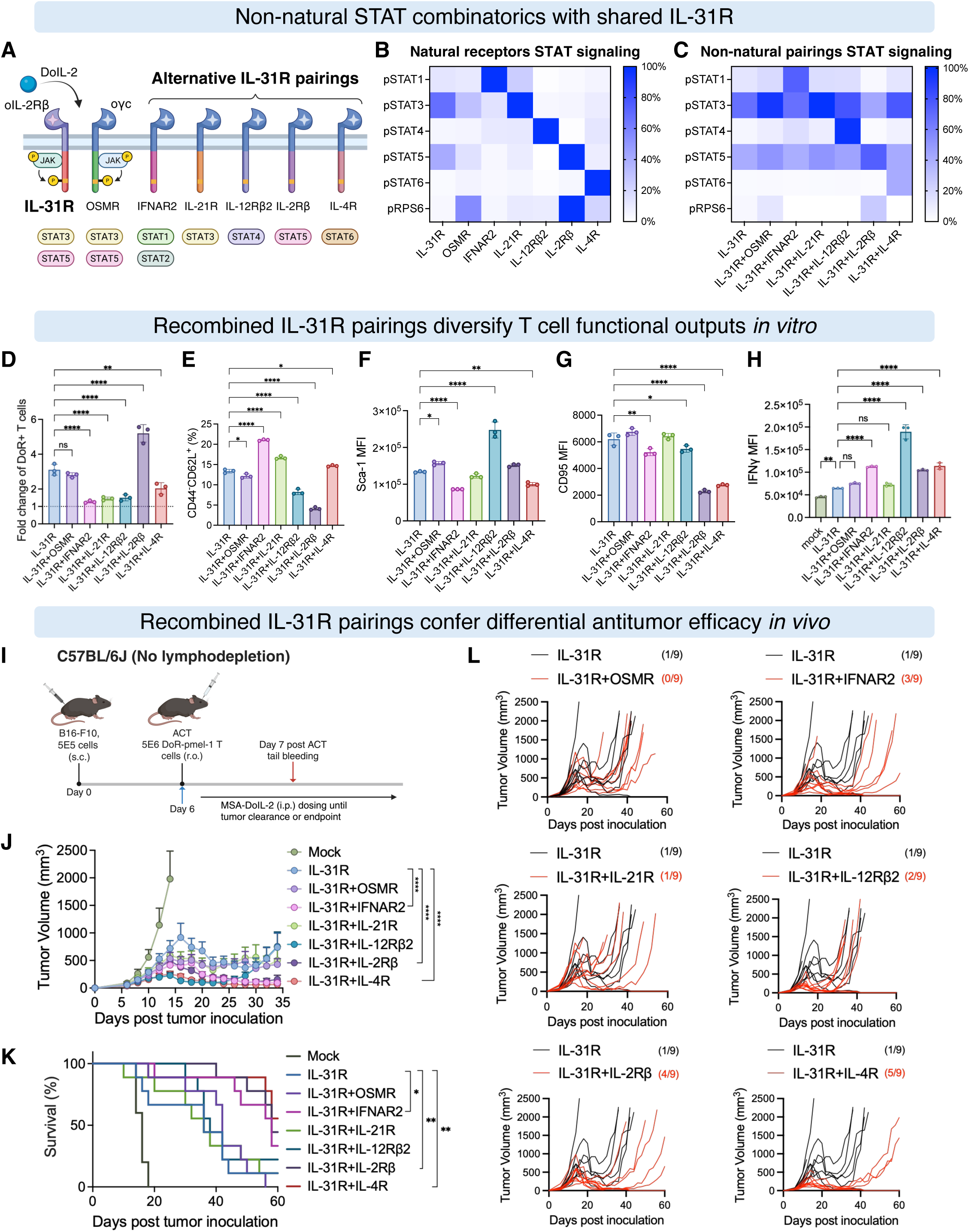
Rewiring a shared IL-31R scaffold with STAT-biased partners reshapes T cell phenotype and antitumor efficacy. (A) Schematic of IL-31R-based receptor pairings. IL-31R, not naturally expressed on T cells, was paired with OSMR (natural partner), IFNAR2, IL-21R, IL-12Rβ2, IL-2Rβ, or IL-4R to generate non-natural receptor heterodimers with distinct STAT activation profiles, as indicated. (B) Heatmap of pSTAT and pRPS6 activities for each individual ICD (IL-31R fused to oIL-2Rβ; partner receptors fused to oγc) activated in isolation by a truncated partner chain containing only the JAK-binding domain. (C) Heatmap of combined pSTAT and pRPS6 activities for each complete IL-31R-based receptor pairing. Data in (B-C) represent technical duplicates; values are normalized maximal MFI from dose-response curves. (D-H) *In vitro* functional characterization of IL-31R-based pairings in Pmel-1 T cells stimulated with DoIL-2 (100 nM, 3 days): proliferative expansion (D); CD44^−^CD62L^+^ Tscm frequency (E); Sca-1 MFI (F); CD95 MFI (G); intracellular IFN-γ MFI (H). Data represent three biological replicates, mean ± SD. (I-L) *In vivo* antitumor efficacy in non-lymphodepleted B16F10 model. Female C57BL/6 mice received ACT of 5×10⁶ Pmel-1 T cells expressing indicated IL-31R-based pairings (day 6), followed by intraperitoneal MSA-DoIL-2 (15 μg, every other day until tumor clearance or endpoint; n = 9 per group). (I) Experimental timeline. (J) Average tumor growth curves. (K) Survival curves. (L) Individual tumor growth curves with complete response (CR) rates indicated. Statistical tests: one-way ANOVA (D-H); two-way ANOVA (J); log-rank Mantel-Cox (K). ns, not significant; *p < 0.05; **p < 0.01; ****p < 0.0001. See also **Figure S5.**

These synthetic combinations yielded distinct and interpretable phenotypes *in vitro*. IL-2Rβ pairing uniquely enhanced proliferation, consistent with STAT5-mediated proliferative function **(Fig. 5D)**; Tscm frequency was highest with IFNAR2 pairing and lowest with IL-2Rβ, while Sca-1 and CD95 each exhibited distinct receptor-dependent expression patterns **(Fig. 5E-G)**; and IFN-γ secretion was greatest with IL-12Rβ2, consistent with STAT4-driven IFN-γ induction, and was modestly elevated by IFNAR2, IL-2Rβ, and IL-4R **(Fig. 5H)**.

We next evaluated the therapeutic consequences of these synthetic STAT combinations in immunocompetent B16F10 tumor-bearing mice (**Fig. 5I**). All pairings were well tolerated (**Fig. S5C**) and delayed tumor growth versus Box control (**Fig. 5J**). IFNAR2, IL-2Rβ, and IL-4R pairings conferred the most durable survival benefit, while OSMR, IL-21R, and IL-12Rβ2 pairings did not significantly extend survival (**Fig. 5K**). Notably, the IL-12Rβ2 pairing achieved the greatest early tumor regression, consistent with its strong IFN-γ production *in vitro*, yet exhibited the lowest abundance of transferred T cells in the peripheral blood, resulting in rapid tumor relapse and no improvement in overall curative responses **(Fig. 5L, Fig. S5D-E)**. These findings indicate that maximizing effector function alone is insufficient for durable tumor control, highlighting the importance of balancing immediate effector activity with long-term T cell persistence.

Collectively, these findings demonstrate that rewiring a common IL-31R scaffold with STAT-biased receptor modules extends signaling landscape beyond the endogenous cytokine receptor repertoire.

### IL-31R pairings drive distinct transcriptional programs that converge on a shared effector core and are concordant with clinical response

To understand why the IL-31R pairings differed in efficacy, we performed single-cell RNA sequencing on tumor-infiltrating Pmel-1 T cells across all conditions, profiling 11,086 TILs and identifying 11 clusters (**Fig. S6A**). Because IL-21R pairing showed functional redundancy with IL-31R alone, our analysis focused on OSMR, IFNAR2, IL-12Rβ2, IL-2Rβ, and IL-4R pairings.

Each receptor pairing predominantly occupied a distinct cluster with a unique transcriptional identity (**Fig. 6A-C**). IL-31R/OSMR closely resembled IL-31R alone, whereas alternative receptor pairings redirected cells into distinct transcriptional states: IFNAR2 pairing preferentially occupied an ISG-enriched C8 cluster; IL-12Rβ2 pairing generated a cluster marked by elevated *Ifng*, *Il18r1*, and *Il18rap* expression; IL-2Rβ pairing preferentially occupied a cytotoxic cluster expressing *Gzmc* and *Gzmf*; and IL-4R pairing generated a distinct cluster enriched for the integrin receptor Itga1 (**Fig. S6B**). Thus, within a shared IL-31R scaffold, changing the partner receptor was sufficient to redirect transcriptional identity.

**Figure 6.**
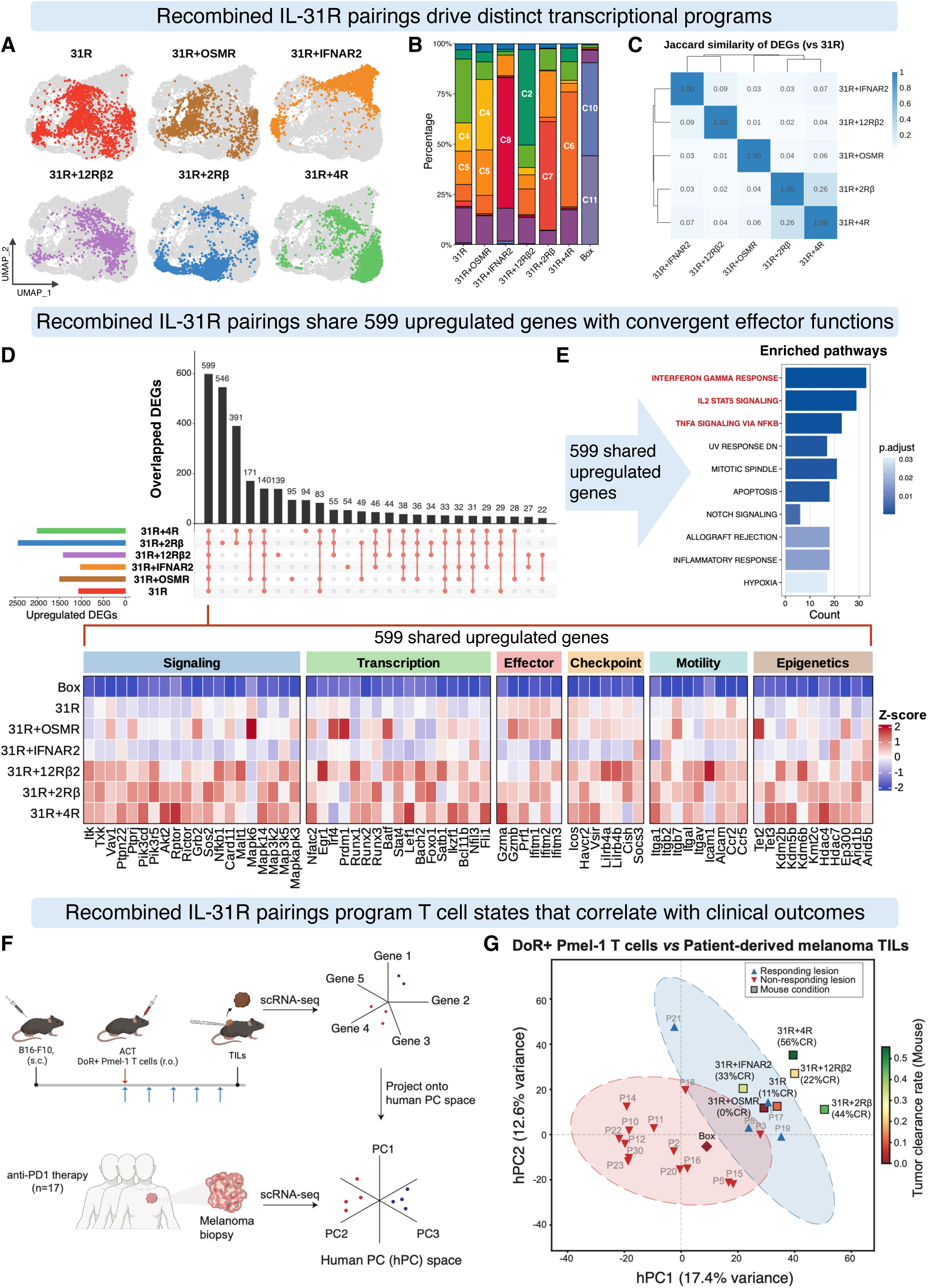
IL-31R-based receptor pairings drive distinct transcriptional programs that converge on a shared effector core and align with clinical response. (A) Per-condition UMAPs showing cell distributions for each IL-31R-based pairing. (B) Stacked bar plots of cluster proportions per sample, with dominant clusters labeled. (C) Heatmap showing pairwise transcriptional similarity among indicated IL-31R-based pairings relative to IL-31R alone. The Jaccard similarity index was used to calculate similarity based on genes with absolute log₂ fold change > 0.5. (D) UpSet plot of upregulated gene overlaps across six IL-31R-based pairings versus Box control (log2FC > 0.5, adjusted P < 0.05). Bar heights indicate intersection sizes; 599 genes are shared across all six conditions. Heatmap of normalized expression (Z-score) for representative genes from the 599-gene shared program, grouped by functional category: signaling, transcription, effector, checkpoint, motility, and epigenetics. (E) Hallmark gene set enrichment of the 599 shared genes. Bar plot shows significantly enriched pathways; color intensity represents adjusted P value. (F) Schematic of the TransComp-R cross-species projection approach. Mouse B16F10 TIL scRNA-seq data are projected onto the principal component (PC) space defined by human melanoma TILs from anti-PD-1-treated patients. (G) Projection of Pmel-1 TIL conditions onto the human PC space (hPC1-hPC2), where hPC1 and hPC2 together separate responding from non-responding human tumor lesions. Mouse conditions are overlaid as projected points colored by complete response rate. See also **Figure S6.**

Despite these private identities, 599 genes were commonly upregulated across all six IL-31R-based pairings relative to Box control (**Fig. 6D)**. These shared genes spanned diverse functional categories, including signal transducers, transcription factors, effector molecules, immune checkpoints, and regulators of cell motility and epigenetic programs (**Fig. 6D, bottom**). This shared program was enriched for Hallmark^52^ pathways associated with IFN-γ response, IL-2-STAT5, and TNFα/NF-κB signaling **(Fig. 6E, Fig. S6G)**. Notably, when IL-31R alone was used as the reference instead of Box control, the shared program largely disappeared **(Fig. S6C-F)**, showing that this conserved effector core derives from the IL-31R module itself, while each private receptor partner adds a distinct program.

Finally, we asked whether these mouse T cell programs bear relevance to human clinical responses. Using TransComp-R, a framework for quantitative cross-species mapping of transcriptomic data^53^(**Fig. 6F**), we projected mouse TIL transcriptomic data onto the principal component space defined by human CD8^+^ TILs from melanoma patients treated with anti-PD-1 checkpoint blockade^54^ in which hPC1 and hPC2 separate responding from non-responding lesions **(Fig. 6G, Fig. S6H)**. Mouse IL-31R pairing conditions projected onto this axis in direct proportion to their *in vivo* complete response rates: IL-31R/IL-4R (56% CR) and IL-31R/IFNAR2 (33% CR) mapped deepest into the responder region, while Box control localized within non-responder space. This cross-species concordance, although preliminary, begins to suggest that synthetic receptor pairings could program T cells into states that might have translational relevance to human antitumor immunity, a prospect that requires further investigation.

### Quantitative STAT biases are decoded into divergent transcription-factor programs

Having linked signaling, transcriptional identity, and therapeutical outcome across both the γc-family and IL-31R pairings, we next asked how these quantitative STAT inputs instigate the transcription factor (TF)-driven regulatory programs that define each T cell state^55,56^. We therefore applied SCENIC analysis^57^, which reconstructs TF regulatory networks by identifying co-expressed gene modules sharing regulatory DNA motifs in their promoters (“regulons”), to an integrated atlas combining all 12 receptor pairing conditions and the Box control (**Fig 7A**; integrated UMAP and cluster distributions in **Fig. S7A-D**).

**Figure 7.**
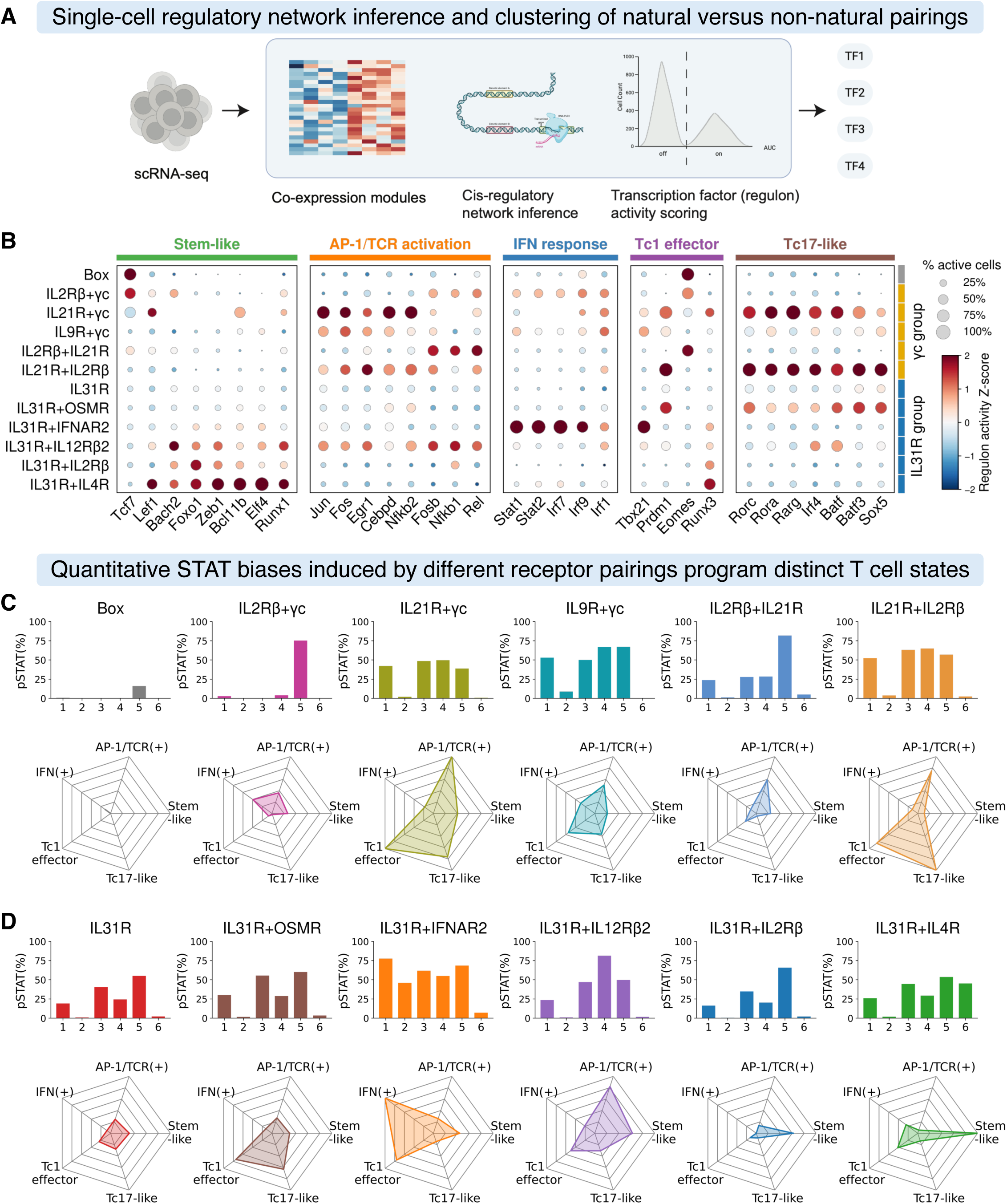
Cytokine receptor rewiring decodes quantitative STAT biases into divergent transcription factor regulatory programs. (A) Schematic of the SCENIC workflow for transcription factor (TF) regulon activity inference. Gene co-expression modules are identified from scRNA-seq data, candidate regulons are defined by cis-regulatory motif enrichment, and regulon activity is scored per cell using AUCell. (B) Dot plot of SCENIC-inferred regulon activities across all receptor pairings. Regulons are selected by cluster-specific regulon specificity scores (RSS) and grouped into five functional axes: stem-like, AP-1/TCR activation, IFN-responsive, Tc1 effector, and Tc17-like. Dot size indicates the fraction of cells with active regulons; color intensity indicates normalized activity (Z-score). (C-D) Integrated analysis of STAT signaling inputs and TF regulatory outputs for γc family pairings (C) and IL-31R-based pairings (D). For each condition: bar plot of relative pSTAT activities derived from Figures 1D and S1B (top); spider plot summarizing relative activity across the five functional regulatory axes (bottom). See also **Figure S7.**

We find that the engineered receptor pairings generated a broad spectrum of regulatory states. Cluster-specific regulon analysis revealed five major functional regulatory axes underlying these states: stem-like (*Tcf7*, *Lef1*, *Bach2*, and *Foxo1*); AP-1/TCR activation (*Jun* and *Fos*); interferon-responsive (*Stat1/2* and *Irf* family); Tc1 effector (*Tbx21*); and Tc17-like (*Rorc*) (**Fig. 7B, Fig. S7E-F**). A pairwise correlation heatmap (**Fig. S7G**) shows these axes are largely mutually exclusive across receptor pairings, though AP-1/TCR activation, Tc1 effector, and Tc17-like programs showed partial co-activation, suggesting the existence of hybrid regulatory states.

For the γc-family pairings, mapping each condition’s STAT input against its regulatory output spider plots illustrates the semblance of a coding logic **(Fig. 7C)**. STAT5-dominant IL-2Rβ/γc maintained a stem-like, *Tcf7*-centered state; adding STAT1/3/4 via the IL-21R arm (IL-2Rβ/IL-21R) shifted cells toward AP-1 activation and early Tc1/Tc17 programs. Strikingly, despite their shared STAT profiles, IL-21R/IL-2Rβ and IL-9R/γc diverged completely at the TF level, IL-21R/IL-2Rβ activating the *Rorc*-centered Tc17 module and IL-9R/γc the *Tbx21*-driven Tc1 module, directly mirroring the lineage divergence of **Fig. 4**.

The IL-31R pairings provided the most controlled test, since all share the IL-31R backbone and differ only in private partner receptor **(Fig. 7D)**. OSMR induced modest Tc17-like activity above IL-31R alone, while each non-natural partner specified a distinct dominant identity: IFNAR2 an interferon-responsive Tc1 state, IL-12Rβ2 an AP-1/TCR-activated state, and IL-2Rβ and IL-4R both a stem-like state built from distinct TF constellations. Notably, IL-31R/IL-4R did not generate the Tc2 state predicted from IL-4R function **(Fig. S7H)**; instead, it produced an emergent stem-like program absent from either receptor alone, an example of a state that cannot be predicted from the parental receptors. Both IL-31R/IL-2Rβ and IL-31R/IL-4R showed the highest stem-like regulatory activity and the most durable antitumor efficacy *in vivo*, directly linking stem-like TF programming to therapeutic benefit.

These analyses establish a linkage map from quantitative STAT input to TF-defined cell state and show that non-natural pairings expand the accessible regulatory landscape beyond the endogenous receptor repertoire.

## Discussion

Evolution has converged on only a small fraction of the possible cytokine receptor pairing space, and in doing so has fixed the vocabulary of JAK-STAT signaling available to immune cells^18,58,59^. A companion study established that across the natural receptor repertoire, quantitative and combinatorial STAT biases, instruct T cell fate, and that the natural receptor landscape constitutes a coarse-grained STAT map whose topology constrains functional divergence^34^. By enforcing pairings across the full all-by-all matrix, we show that this natural vocabulary can be expanded into a much larger synthetic, fine-grained one. Non-natural pairings access emergent T cell states whose identities cannot be read off from either parental receptor. We consider what this expanded landscape implies for how cytokine signaling encodes cell fate, and how that code can be engineered.

A first conceptual implication is that receptor complex geometry is itself an axis of signaling information and diversification, independent of receptor composition, as has been previously suggested in several systems^60–62^. The starkly different behavior of the two IL-2Rβ/IL-21R orientations shows that the same two chains can specify different fates depending simply on which occupies which position in the dimer. Since it is generally established from structural studies that the receptor at site 1 binds to cytokine first, followed by receptor 2 to form the dimer^42,63^, this directionality likely reflects asymmetries in receptor-proximal kinase activity, STAT docking geometry, chain abundance, or ligand-binding orientation, and it effectively doubles the engineerable space into two distinct, orientation-defined coordinates. Natural cytokine complexes, constrained to fixed geometries, cannot exploit this degree of freedom, whereas synthetic systems can, suggesting that orientation is an underused design variable for tuning signaling output.

A second implication concerns the nature of combinatorial signaling itself. One expectation might be that forcing two receptors together would yield an additive combination of their signaling programs, but our data contradict this notion. IL-21R/IL-2Rβ and IL-9R/γc converge on similar STAT1/3/4/5 activation *in vitro* yet program distinct lineages, Tc17-like versus Tc1, *in vivo*. What distinguishes them is not which STATs are engaged but the quantitative ratio among them: a modest excess of STAT1 and STAT4 is critical to the fate decision. This argues that T cell fate is encoded in a continuous, ratiometric STAT space rather than a discrete, identity-based one, and that emergence, non-additive outcomes, is possible. The IL-31R/IL-4R result makes the same point from the opposite direction: a receptor that specifies Tc2 in its natural context instead produced a stem-like program on a synthetic scaffold, a state neither parent encodes alone.

These observations suggest a distinct use for engineered cytokine receptors. Rather than mimicking or amplifying a natural signal, non-natural pairings can position a T cell at coordinates in STAT space not occupied by any natural ligand, accessing states absent from the natural repertoire, whether because they were never selected for or because they are incompatible with normal homeostasis. The fine-grained atlas we describe links signaling coordinates to functional readouts, which could support a more systematic approach to selecting receptor pairings for a desired T cell state.

Our *in vivo* data also clarify an issue of practical importance for cell therapy: proliferative capacity and therapeutic efficacy are separable properties. The canonical IL-2Rβ/γc pairing drove the strongest expansion yet cured no mice, whereas pairings that preserved stem-like or hybrid programs cured despite weaker expansion and even without lymphodepleting preconditioning. The IL-12Rβ2 pairing illustrates the converse failure mode: maximal early effector output followed by rapid collapse and relapse. Durable control therefore depends on the balance a receptor strikes between effector function and persistence, a balance that the atlas allows one to tune by choice of pairing. That a Tc17-like state with retained cytotoxicity could be installed by receptor engineering alone, which is associated with superior persistence, and durable antitumor responses in adoptive cell therapy settings^64–67^, without exogenous polarizing cytokines, points to a general route for building therapeutically favorable states directly into engineered cells.

From a translational standpoint, the all-by-all STAT matrix is a design resource for next-generation adoptive cell therapy: several of the states it encodes, such as Tc17-like, ISG-driven Tc1, and stem-like programs, are exactly the states associated with durable antitumor immunity, and the concordance we observe between synthetic mouse programs and human checkpoint-response signatures suggests these coordinates are conserved. More broadly, the framework is not limited to the receptors or cell type studied here; the same logic could be applied to CD4⁺ subsets, other immune lineages, and additional receptor modules to chart still larger regions of the programmable signaling space.

In sum, the natural cytokine receptor repertoire is a coarse-grained map that can be refined into a fine-grained atlas by sampling pairings evolution left unexplored. This atlas reveals that cytokine signaling encodes fate ratiometrically, that receptor orientation is an independent diversification axis, and that non-natural pairings generate genuinely emergent states, with several conferring superior antitumor efficacy. Systematic exploration of this space offers a strategy for engineering immune cell identities beyond the reach of natural cytokine signaling.

### Limitations of the study

Several limitations should also be considered. First, although we observed pronounced rewired signaling and emergent T cell states, the underlying mechanistic basis of these effects remains incompletely defined. Our conclusions regarding STAT balance are primarily correlative and require direct perturbation-based validation to establish causality. Second, our analyses were restricted to Pmel-1 CD8⁺ T cells, and whether similar principles apply to CD4⁺ T cell subsets or other immune cell types remains to be determined. Furthermore, our *in vivo* findings were primarily derived from the B16F10/Pmel-1 model, and broader validation across diverse tumor models and human systems will be required to establish translational relevance. Finally, only a subset of possible receptor pairings was functionally characterized in this study, leaving much of the programmable signaling space unexplored.

## Acknowledgments

This work was supported by NIH-RO1-AI51321 (K.C.G.), Parker Institute for Cancer Immunotherapy (K.C.G.), Stanford-UCSF Weill Cancer Research Fund, Ludwig Institute, K.C.G. is an Investigator of the Howard Hughes Medical Institute. This work was in part supported by the NIH/NCI 2P01CA049605 (Z.G.), NIH/OD 1OT2OD038101 (Z.G.), Stanford Center for Digital Health Award (Z.G.), Laude Institute MOONSHOTS \\ ONE Honorable Mention Award (Z.G.). Z.G. was supported by the Parker Bridge fellowship (PICI C-02895) and the NIH/NCI Pathway to Independence award (1K99CA293149, 4R00CA293149). K.C.G. and Z.G. are members of the Parker Institute for Cancer Immunotherapy, which supports the Stanford University Cancer Immunotherapy Program. K.C.G. and Z.G. are Stanford-UCSF Weill Cancer Hub West Investigators. The content is solely the responsibility of the authors and does not necessarily represent the official views of the foundations and the NIH. We acknowledge the Animal care facilities at Stanford, Stanford Shared FACS Facility for their technical assistance. Schematics and diagrams were created using BioRender.

## Author contributions

K.C.G. conceived the study, supervised the experiments, and collaborated with the other authors on the manuscript. P.T. conceived several aspects of the study, designed and conducted the experiments, analyzed the data, and wrote the manuscript. K.T. and Z.G. contributed to the analysis of scRNA-seq data. Y.Z. and H.J. assisted with *in vivo* mouse experiments. All authors contributed to manuscript preparation.

## Data availability

The raw and processed scRNA-seq data generated in this study will be deposited for public availability in the GEO database upon publication. The code used for RNA-seq data analysis will also be shared publicly at that time. Any additional information required to reanalyze the data reported in this paper is available from the lead contact upon request.

## Methods

### Experimental model and study participant details

#### Mice

Six- to eight-week-old Thy1.2^+^ C57BL/6 (C57BL/6J) mice were purchased from Jackson Laboratory. Thy1.1^+^ pmel-1 TCR-transgenic mice (B6.Cg-Thy1a/Cy Tg(TcraTcrb)8Rest/J) and NSG (NOD.Cg-Prkdcscid Il2rgtm1Wjl/SzJ) mice were originally purchased from the Jackson Laboratory and maintained in the Stanford University-Lorry Lokey (SIM1) Facility. All animal experiments were conducted in accordance with the guidelines of the protocol (ID: 32279) approved by the Institutional Animal Care and Use Committee (IACUC), Administrative Panel on Laboratory Animal Care (APLAC) at the Stanford University.

#### Cell lines and cell culture

The B16-F10 mouse melanoma cell line and A375 cell was purchased from ATCC and cultured with complete DMEM medium (high-glucose DMEM supplemented with 10% FBS, 2 mM GlutaMAX, 10 mM HEPES, and penicillin-streptomycin). Platinum-E (Plat-E) and Platinum-GP (Plat-GP) retroviral packaging cell lines were purchased from Cell Biolabs and cultured with complete DMEM medium. Expi293F cells were purchased from Gibco and cultured in suspension with Expi293 Expression Medium. All related cell culture reagents were purchased from Gibco.

#### Primary T cells preparation and culture

Primary mouse total T cells or CD8^+^ T cells were isolated from spleen and lymph nodes of six- to eight-week-old mice using Pan T Cell Isolation Kit II (Miltenyi) or CD8a^+^ T Cell Isolation Kit (Miltenyi) and cultured in complete RPMI 1640 medium (RPMI 1640 supplemented with 10% FBS, 50 μM 2-mercaptoethanol, non-essential amino acids, sodium pyruvate, 2 mM GlutaMAX, 10 mM HEPES, and penicillin-streptomycin) supplemented with 100 U/mL mouse IL-2 (Miltenyi). Human T cells were isolated from thawed PBMC using EasySep™ Human T Cell Isolation Kit (STEMCELL) and cultured in complete RPMI 1640 medium supplemented with 100 U/mL human IL-2 (PeproTech). Human PBMC were isolated from LRS chambers (Stanford Blood Center) and cryopreserved until time of use.

### Method details

#### Protein production

DNA encoding mouse and human double-orthogonal IL-2 (DoIL-2) were cloned into the mammalian expression vector pD649, which includes a C-terminal 6x His tag for affinity purification. DNA encoding mouse serum albumin (MSA) was cloned into the N-terminal of the pD649 constructs described above to express MSA fusion protein. Mammalian expression DNA constructs were transfected into Expi293F cells using the Expi293 Expression System (Gibco) for secretion and purified from the clarified supernatant by nickel affinity resin (Ni-IMAC, Thermo Fisher Scientific) followed by size-exclusion chromatography with a Superdex-200 column (Cytiva) and formulated in sterile phosphate-buffered saline (PBS). Endotoxin was removed using the Proteus NoEndo HC Spin column kit (VivaProducts) and endotoxin removal was confirmed using the Pierce LAL Chromogenic Endotoxin Quantification Kit (Thermo Fisher Scientific). Proteins were concentrated, frozen with liquid nitrogen and stored at -80°C until use.

#### Retroviral vectors for mammalian expression

To characterize the signaling matrix of all-by-all receptor pairs, cDNA encoding each mouse chimeric cytokine receptor fused with the orthogonal human IL-2Rβ ectodomain was cloned into the retroviral pMSCV vectors containing TagBFP by PCR and isothermal assembly (ITA). Meanwhile, cDNA encoding each mouse chimeric cytokine receptor fused with the orthogonal human γc ectodomain was cloned into the retroviral pMSCV vectors containing ZsGreen. To coexpress two orthogonal cytokine receptors in one vector, a P2A self-cleaving peptide was employed to enable the bicistronic expression of both orthogonal receptors at equal level. Box regions of all natural cytokine receptors were defined by UniProt and related references^38,39^. Geneblock (IDT) encoding NY-ESO-1 specific TCR (1G4-α95:LY) was cloned into the retroviral pMSCV vectors containing EYFP.

#### Retrovirus production

To generate retrovirus for the transduction of mouse T cells, Plat-E cells (Cell Biolabs) were seeded at 8 × 10^5^ cells per well with 3 mL medium in a 6-well tissue culture treated plate. Following an overnight incubation, fresh media was replenished prior to transfection. For each well of transfection, 5 µg of plasmid (3 µg of pMSCV retroviral expression vector plus 2 µg of pCL-Eco packaging vector) was added to 300 µl of Opti-MEM I Reduced Serum Medium (Gibco), followed by 15 µl of FuGENE® HD transfection reagent (Promega), with gentle vortexing. After a 15-minute incubation, the transfection mixture was gently added to Plat-E cells. Retrovirus supernatant was collected at 48 hours post-transfection and filtered through a 0.45-µm syringe filter (Nalgene). If not used immediately, the virus can be stored at 4 °C for 1 week or frozen at - 80°C for 6 months.

To generate retrovirus for the transduction of human T cells, Plat-GP cells (Cell Biolabs) were prepared under the same condition but virus production was using a different plasmid transfection recipe (3 µg of pMSCV retroviral expression vector plus 2 µg of pLTR-RD114A envelope plasmid). Retrovirus supernatant was collected at 48 hours post-transfection and filtered through a 0.45-µm syringe filter (Nalgene). If not used immediately, the virus can be stored at 4 °C for 1 week or frozen at -80°C for 6 months.

#### Activation and transduction of primary T cells

Isolated T cells from C57BL/6 mouse were activated with plate-bound anti-mouse CD3ε (5 μg/mL, clone 145-2C11, Biolegend) and soluble anti-mouse CD28 (5 μg/mL, clone 37.51, Biolegend) in complete RPMI 1640 medium supplemented with 100 U/mL mouse IL-2 (Miltenyi) for 24 h. Isolated CD8+ T cells from pmel-1 mouse were activated with 1 µM human gp100 peptide (GenScript) in complete RPMI 1640 medium supplemented with 100 U/mL mouse IL-2 (Miltenyi) for 24 h. One day before transduction, 6-well tissue culture plates were coated with Retronectin (25 μg/mL, Takara) and placed in a 4 °C refrigerator overnight. The following day, plates were washed twice with complete RPMI 1640 medium. Viral supernatant (3 mL) was added to each well and spun at 2500 g for 2 hours at 32°C. After spinning, the virus particles were captured on the plate. Activated T cells (2 × 10^6^ in 3 mL medium) were added to each well after aspirating the viral supernatant and spun at 1000 g for 10 minutes at 32°C. After incubating overnight, T cells were collected and expanded in a fresh medium until further analysis. On day 3, transduction efficiency was assessed based on the expression of fluorescent protein using flow cytometry. Untransduced T cells activated and cultured in parallel were used as control. T cells were collected for *in vitro* assays or *in vivo* injection three days after spinfection.

Isolated human T cells were activated with plate-bound anti-human CD3ε (5 μg/mL, clone OKT-3, Biolegend) and soluble anti-human CD28 (5 μg/mL, clone CD28.2, Biolegend) for 48 h. Activated T cells were collected and transduced with viral supernatant on 6-well plates coated with Retronectin as described above. To co-transduce T cells with two different retroviruses, a secondary transduction was performed 12 hours later after the first spinfection. Transfected human T cells were collected and expanded in a fresh medium until they returned to a resting state for functional assays. For *in vivo* injection, human T cells were expanded and collected for 10-12 days after activation.

#### Phosphoflow signaling assays of primary T cells

Actively growing mouse or human primary T cells were washed and rested in T cell medium without IL-2 for 12-18 hours before signaling assays. Cells were plated in a 96-well round bottom plate in complete RPMI 1640 medium without FBS prior to the assay. To detect phosphorylation of STAT proteins, Ribosomal Protein S6 (RPS6), T cells were stimulated by addition of orthogonal cytokines for 30 minutes at 37°C. To detect phosphorylation of Erk, p38 MAPK, Akt and p90RSK, T cells were stimulated by addition of orthogonal cytokines for 5 minutes at 37°C. The reaction was terminated by fixation with 2.1% paraformaldehyde (BD Cytofix) for 30 minutes at 37°C. Fixed cells were washed and permeabilized with ice-cold methanol (BD Phosflow Perm Buffer III) for 30 minutes on ice or stored at -80°C for later analysis. Cells were washed with staining buffer (PBS containing 2% FBS) before staining with phosphoflow antibodies (BD or CST) for 30 min to 1 hour at 4°C in the dark. Cells were washed and analyzed on a CytoFlex (Beckman Coulter) or Novocyte Quanteon (Agilent Technologies). For the specific pSTAT2 detection in mouse T cells, ectopically transduced human STAT2 was used as a surrogate, and analyzed under the positive gate. Data represent the median fluorescence intensity (MFI), and points were fit to the [agonist] versus dose–response (three parameters) model using Prism 10 (GraphPad).

#### Mouse T cell *in vitro* proliferation and immunophenotyping

Actively growing transduced mouse T cells were washed and re-suspended in mouse T cell medium lacking IL-2 and seeded at a density of 50,000 T cells per well (in 100 µL) in a 96-well round bottom tissue culture plate. T cell proliferation was stimulated by addition of 100 µL DoIL-2 (200 nM) to a total volume of 200 µL. Before the culture, 100 µL T cells were transferred to a new plate for cell counting as day 0. The remaining 100 µL T cells were cultured for 3 days at 37 °C. On day 3, T cells were collected and counted by FACS using the CytoFLEX equipped with a high throughput sampler as compared with day 0. The total number of live orthogonal cytokine receptor-expressing T cells were gated by staining of Fixable Viability Dye eFluor™ 780 (eBioscience) and expression of mCherry. For the immunophenotyping of T cell stemness, CD44, CD62L, Sca-1 and CD95 antibodies were included in the staining buffer and gated during cell counting.

#### Mouse T cell *in vitro* differentiation and immunophenotyping

Actively growing transduced mouse T cells were washed and re-suspended in mouse T cell medium lacking IL-2 and seeded at a density of 5× 10^5^ T cells per well (in 0.5 mL) in a 24-well tissue culture plate. T cells were first stimulated by addition of 0.5 mL medium containing DoIL-2 (200 nM) and human gp100 (200nM) to a total volume of 1 mL. After 72 hours of stimulation, T cells were collected and seeded into the 96-well round bottom tissue culture plate. For the staining of lineage specific transcription factors, T cells were first stained by Fixable Viability Dye eFluor™ 780 (eBioscience) and intracellular staining was subsequently performed using the True-Nuclear Transcription Factor Buffer Set (BioLegend). For the staining of indicated secreted cytokines, Cell Stimulation Cocktail plus Protein Transport Inhibitor (eBioscience) was added into the T cell medium for 4-6 hours incubation at 37 °C. After cell viability staining, intracellular cytokines staining was subsequently processed using the Cell Fixation/Permeabilization Kit (BD Biosciences).

#### *In vivo* syngeneic mouse tumor model

Six to eight-week-old C57BL/6J mice were subcutaneously injected with 5 × 10^5^ B16F10 tumor cells in 100 μL of PBS into the right flank. On day 6, mice were randomized based on average tumor size and received adoptive transfer of 5 × 10^6^ pmel T cells (50% transduced). T cells were resuspended in 50 μl of PBS per mouse and administered by retroorbital intravenous injection. MSA-mDoIL-2 (15 μg) in 100μl of PBS was administered the same day and every other day until day 20. Peripheral blood (10 μL) was collected using Microhematocrit Capillary Tubes (Fisherbrand) at indicated time points from the tail vein for quantification of adoptively transferred pmel T cells by flow cytometry. Tumor size (length and width) was measured with calipers three times a week and volume was calculated as (length × width^2^)/2. Post-therapy survival of mice was monitored for at least 60 days post-tumor inoculation. Mice were euthanized when the total tumor volume exceeded 2,000 mm^3^ or reached the morbidity criteria, as per APLAC guidelines.

#### *In vivo* human tumor model in immunodeficient NSG mice

Six to eight-week-old NSG mice were subcutaneously injected with 1 × 10^6^ A375 tumor cells in 100 μL of PBS into the right flank. On day 8, mice were randomized based on average tumor size and received adoptive transfer of 5 × 10^6^ human T cells (50%-60% transduced with both NY-ESO-1 TCR and orthogonal receptors). T cells were resuspended in 50 μl of PBS per mouse and administered by retroorbital intravenous injection. MSA-DoIL-2 (15 μg) in 100μl of PBS was administered the same day and every other day until day 20. Peripheral blood (10 μL) was collected using Microhematocrit Capillary Tubes (Fisherbrand) at indicated time points from the tail vein for quantification of adoptively transferred human T cells by flow cytometry. Tumor size (length and width) was measured with calipers three times a week and volume was calculated as (length × width^2^)/2. Post-therapy survival of mice was monitored for at least 60 days post-tumor inoculation. Mice were euthanized when the total tumor volume exceeded 2,000 mm^3^ or reached the morbidity criteria, as per APLAC guidelines.

#### Single-cell RNA sequencing of *in vitro*-stimulated mouse T cells

CD8 T cells isolated from pmel-1 mice were activated and transduced with indicated double-orthogonal cytokine receptors individually. Engineered T cells (mCherry^+^) were sorted at 48 hours post transfection. After an overnight resting, T cells were stimulated with 100 nM DoIL-2 for 24 hours to initiate the transcription programs. After the stimulation, live T cells were purified with the Dead Cell Removal Kit (Miltenyi) and fixed using the Evercode Cell Fixation v3 Kit (Parse Biosciences). Fixed cells underwent three rounds of combinatorial barcoding according to the protocol of Evercode Cell WT Mini v3 Kit (Parse Biosciences) and resulted in 2 sublibraries for sequencing. The generated 3′ Gene Expression libraries were sequenced using the NovaSeq X Plus (illumina) with a sequencing depth of > 50K paired-end reads per cell.

#### Single-cell RNA sequencing of tumor infiltrating mouse T cells

C57BL/6 mice were subcutaneously inoculated with B16F10 tumor cells (1 × 10^6^) into the right flank. On day 7, tumor-bearing mice were randomized and received the adoptive transfer of 5 × 10^6^ engineered pmel T cells by retroorbital intravenous injection and followed by intraperitoneal administration of MSA-mDoIL2 (15 μg) every other day for a total of 5 times. On day 17, mice were sacrificed, tumors were minced and dissociated using a mouse tumor dissociation kit (Miltenyi Biotec) and a gentleMACS Octo Dissociator (Miltenyi Biotec). Tumor-infiltrating pmel T cells were first enriched using mouse CD45 (TIL) MicroBeads (Miltenyi Biotec), and then sorted with surface markers (CD8, Thy1.1), viability dye (eBioscience) and expressed fluorescent protein. Sorted T cells were subjected to fixation using the Evercode Cell Fixation v3 Kit (Parse Biosciences) and combinatorial barcoding using Evercode Cell WT Mini v3 Kit (Parse Biosciences) to generate 2 sublibraries for sequencing. The sequencing parameters and sample demultiplexing process were described as above.

#### Analysis of single-cell RNA sequencing data

The raw FASTQ files from sub-libraries were demultiplexed into all individual samples using the pipeline from Trailmaker pipeline (Parse Biosciences). Reads were aligned to the mouse reference genome (GRCm39), and gene annotations were based on Ensembl release 109. Gene-level count matrices were generated using default parameters. Initial quality control, including filtering of low-quality cells based on UMI counts, mitochondrial gene content, and doublet detection, was performed using the built-in automated pipeline in Trailmaker (Parse Biosciences). Gene expression matrices were log-normalized, and 2,000 highly variable genes (HVG) were identified. Principal component analysis (PCA) was performed, and the top 30 principal components were used for neighborhood graph construction. Cells were clustered using the Leiden algorithm, and low-dimensional visualization was performed using Uniform Manifold Approximation and Projection (UMAP). Downstream analyses and visualization were performed using the Seurat v5 package in R and Scanpy (v1.10.3) in Python.

Differential gene expression analysis was performed using the Wilcoxon rank-sum test implemented in Seurat. Genes detected in at least 10% of cells in either group were considered for testing. Differentially expressed genes (DEGs) were defined as those with an adjusted P value < 0.05 after Benjamini-Hochberg correction and an absolute log2 fold change greater than 0.5. Cluster marker genes, receptor-specific transcriptional signatures, and pairwise comparisons between selected cell populations were identified using this framework. DEGs were subsequently used for cluster annotation, functional interpretation, and downstream pathway enrichment analyses. Functional enrichment analysis of DEGs was performed using clusterProfiler. Gene Ontology (GO) biological process and Kyoto Encyclopedia of Genes and Genomes (KEGG) pathway analyses were conducted using upregulated genes unless otherwise specified. Gene set enrichment analysis (GSEA) was performed using ranked gene lists ordered by log2 fold change. For transcriptional regulatory network analysis, regulon activity was inferred using pySCENIC (v0.12.1). Briefly, gene co-expression networks were constructed using GRNBoost2, and candidate regulons were identified with RcisTarget based on motif enrichment analysis. RcisTarget motif rankings based on the mm10 reference databases (500 bp upstream and 10 kb around TSS) were used. Regulon activity in individual cells was then quantified using AUCell, generating area under the curve (AUC) scores at the single-cell level. Regulon activity scores were used to define transcription factors-driven cell states, and regulon specificity scores (RSS) were calculated to identify cluster-specific transcription factors. Scaled regulon activity was used for visualization. Pairwise Pearson correlations were computed between selected regulons to identify co-activity relationships among transcription factor regulons.

To evaluate the clinical translatability of engineered cytokine receptor conditions, we adapted the TransComp-R framework to project mouse 31R Pmel-1 TIL conditions into a principal component space defined by human anti-PD-1-treated CD8^+^ T cells. Smart-seq2 profiles of post-treatment CD45^+^ tumor-infiltrating immune cells were obtained from GSE120575^41^. Response was annotated at the lesion level (responder, R; non-responder, NR). CD8 T cells were identified by marker-based gating (CD8A/CD8B TPM > 1 with CD3D/CD3E co-expression, excluding CD14/CD68/CD163 and MS4A1/CD19), myeloid fractions were removed. TPM matrix was filtered to remove clonotype (TR/IG-V/J), sex-linked, mitochondrial, ribosomal, housekeeping, and unannotated genes, as well as genes detected in <5% of cells. Lesion-level pseudobulk profiles were computed as the mean TPM across CD8 T cells per lesion (≥ 10 cells; n = 20 lesions, 4 R / 16 NR) and log2 transformed. One-to-one mouse-human orthologs (biomaRt/Ensembl) were used to align features, and the top 2,000 highly variable genes (by coefficient of variation across mouse 31R conditions) were retained. Expression was rank-transformed within each sample and gene-wise z-scored using a StandardScaler fit on the human data; PCA was performed on the human matrix (scikit-learn) retaining min(n-1, 10) components, and mouse condition-level pseudobulks (similarly rank- and z-transformed with the human-fitted scaler) were projected into this space via the human loadings. Human lesions are shown as points with 95% confidence ellipses for R and NR; mouse 31R conditions are overlaid as projected points colored by complete-response rate.

#### Statistical analysis

Statistical analysis was performed using GraphPad Prism v10, except where indicated. All values and error bars are shown as mean ± SEM. Comparisons of two groups were performed by using two-tailed unpaired Student’s t test. Comparisons of multiple groups were performed by using one-way analysis of variance (ANOVA) with Tukey’s multiple-comparisons test unless otherwise indicated. Experiments that involved repeated measures over a time course, such as tumor growth were performed by using two-way ANOVA with Tukey’s multiple-comparisons post-test. Survival data were analyzed using the Log-rank (Mantel-Cox) test. P-values were considered significant if less than 0.05, indicated as ∗ p <0.05, ∗∗ p <0.01, ∗∗∗ p <0.001, and ∗∗∗∗ p <0.0001. No statistically significant (NS) differences were considered when P-values were larger than 0.05.

## Supplement information

Table S1. Amino acid sequences of double-orthogonal cytokine receptors.

Table S2. Differential gene expression analyses of *in vitro*-stimulated T cells expressing γc family receptor pairings.

Table S3. Differential gene expression analyses of TILs expressing γc family receptor pairings.

Table S4. Differential gene expression analyses of TILs expressing IL-31R-based receptor pairings.

Table S5. Cluster-specific transcription factor regulons ranked by regulon specificity score (RSS) in integrated TIL scRNA-seq data.

## Declaration of generative AI and AI-assisted technologies in the writing process

During the preparation of this work, the authors used Claude and ChatGPT to improve the readability of the manuscript and to assist with the analysis and visualization of single-cell RNA sequencing data. The authors reviewed and edited the output as needed and take full responsibility for the content of the published article.

## Supplemental Figures and Legends

**Figure S1.**
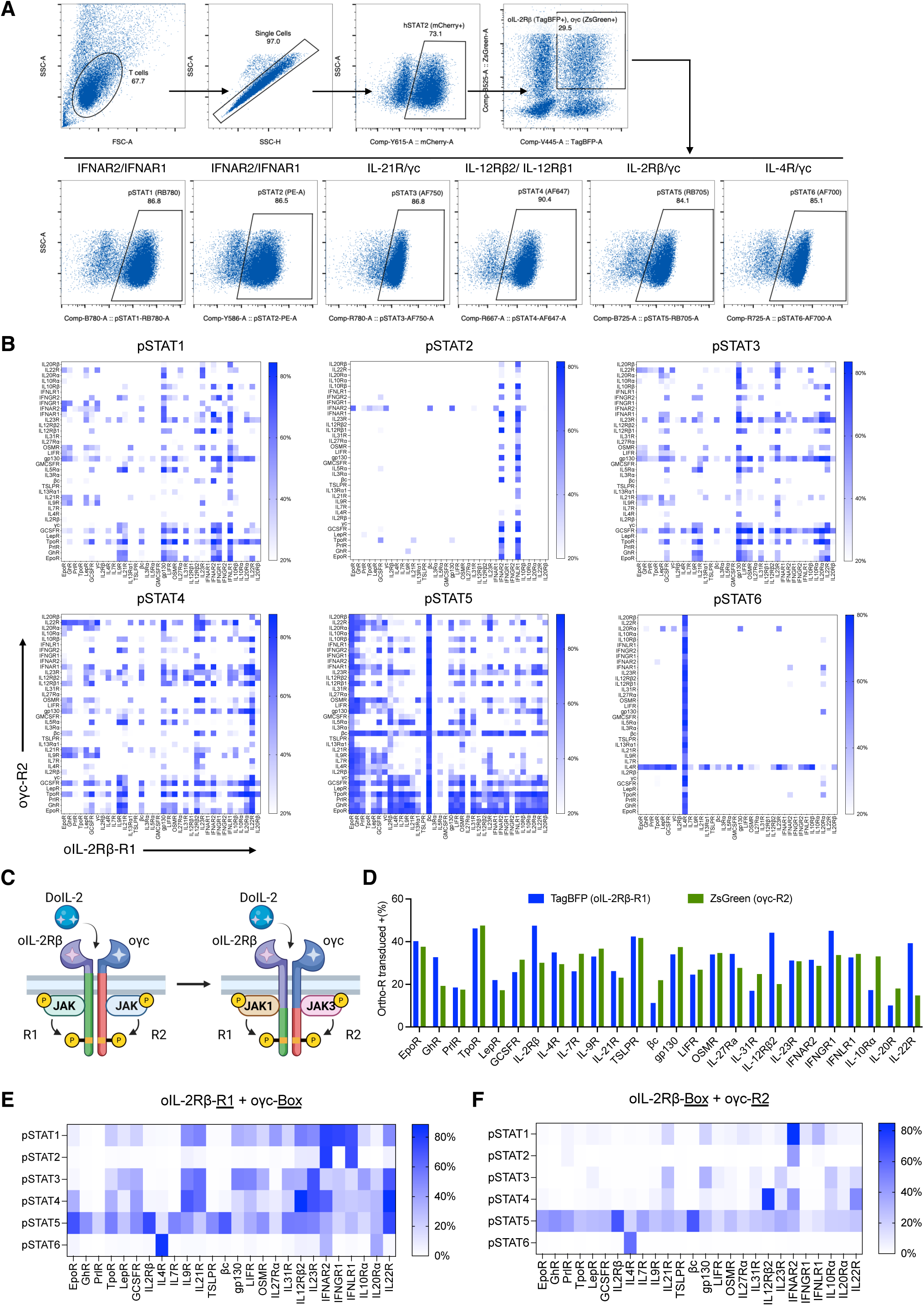
Engineering double-orthogonal cytokine receptors and systematic profiling of pSTAT signaling, related to Figure 1. (A) Flow cytometry gating strategy to identify double-orthogonal cytokine receptors-transduced T cells and quantify pSTAT-positive populations across STAT1-STAT6. mCherry-tagged human STAT2 was expressed to enable STAT2 activity detection. (B) Heatmap representation of the pSTAT signaling interactome resulting from all pairwise combinations of natural cytokine receptor ICDs with intact JAK- and STAT-binding regions (36 × 36). (C) Design of orthogonal chimeric cytokine receptors by replacing the native ICDs with engineered ICDs containing only STAT-binding regions while preserving a fixed JAK1/JAK3 pair. (D) Expression validation of orthogonal chimeric cytokine receptors with engineered ICDs, including cytokine receptors containing putative STAT-binding regions. (E) Heatmap of pSTAT signaling activity induced by engineered intracellular domains fused to oIL-2Rβ and paired with truncated oγc containing only the JAK3-binding region. (F) Heatmap of pSTAT signaling activity induced by engineered intracellular domains fused to oγc and paired with truncated oIL-2Rβ containing only the JAK1-binding region.

**Figure S2.**
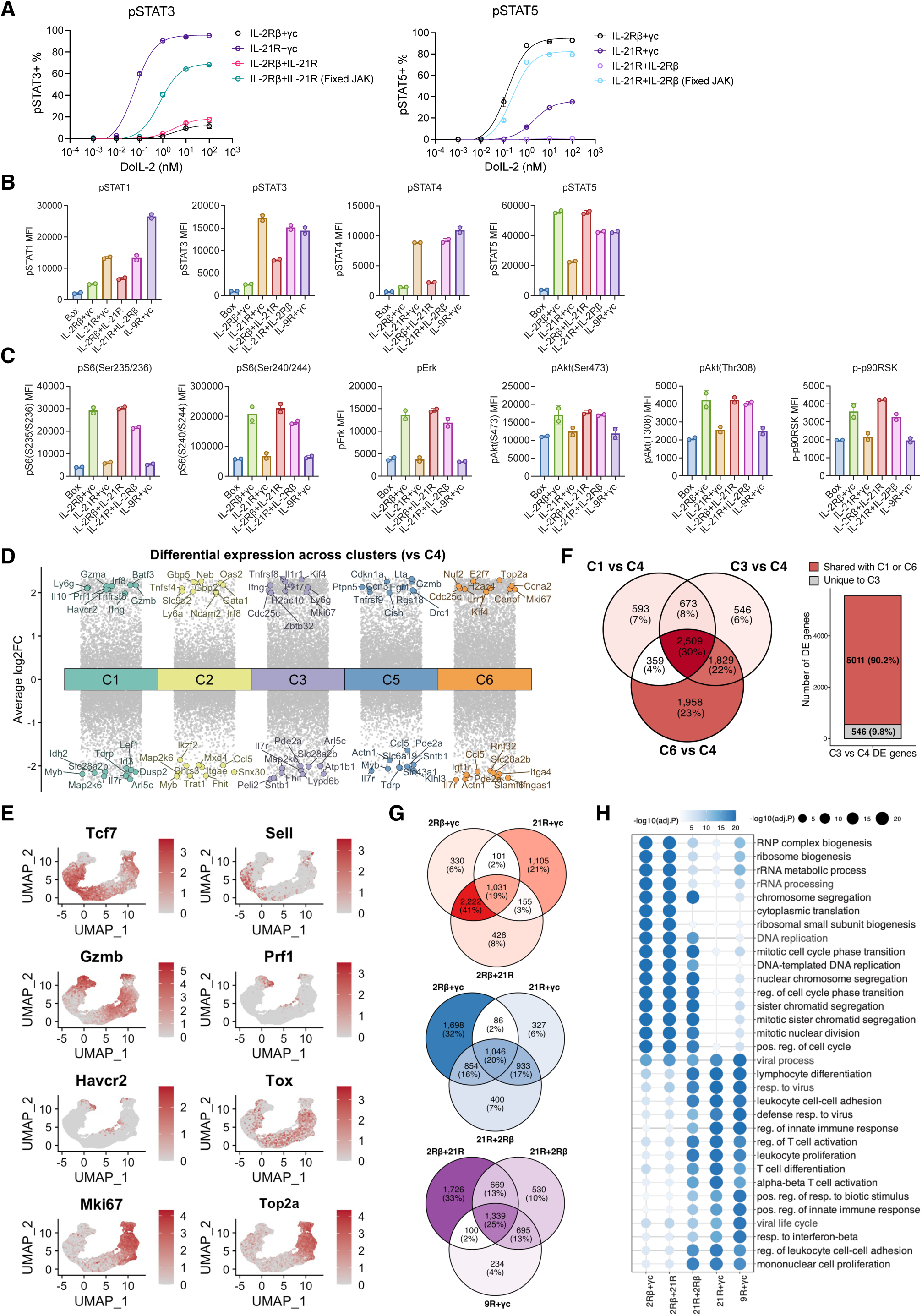
Signaling and transcriptional characterization of rewired IL-2Rβ–IL-21R heterodimers and corresponding natural receptor pairings, related to Figure 2. (A) Dose-response curves of pSTAT3 and pSTAT5 signaling for natural γc receptor pairings (IL-2Rβ/γc and IL-21R/γc) and unnatural IL-2Rβ–IL-21R heterodimers with distinct orientations and JAK-binding domains. (B-C) Bar plots showing median fluorescence intensity (MFI) of intracellular phosphoprotein staining in the indicated T cells after stimulation for STAT (B) and non-STAT (C) signaling, related to Figure 2B. Data represent technical duplicates. (D) Differential gene expression across annotated clusters relative to C4, with each gene plotted by average log₂ fold change (log₂FC). Selected cluster-enriched genes are highlighted and labeled. (E) UMAP visualization of gene expression for selected markers associated with T cell functional states, related to Figure 2J. Color intensity indicates normalized expression levels. (F) Overlap of differentially expressed genes (DEGs) across C1 vs C4, C3 vs C4, and C6 vs C4 comparisons. Venn diagram shows shared and unique DEGs among clusters. Bar plot summarizes the proportion of C3 vs C4 DEGs that are shared with C1 or C6 versus those unique to C3. Numbers indicate gene counts and percentages. (G) Venn diagrams showing overlap of differentially expressed genes among unnatural IL-2Rβ/IL-21R heterodimers and natural γc receptor pairings (IL-2Rβ/γc, IL-21R/γc and IL-9R/γc). Numbers indicate gene counts and percentages. (H) GO biological process enrichment of upregulated genes for each receptor pairing versus Box control (adjusted P < 0.05, log2FC > 0.5). The union of the top 10 enriched terms per condition is shown.

**Figure S3.**
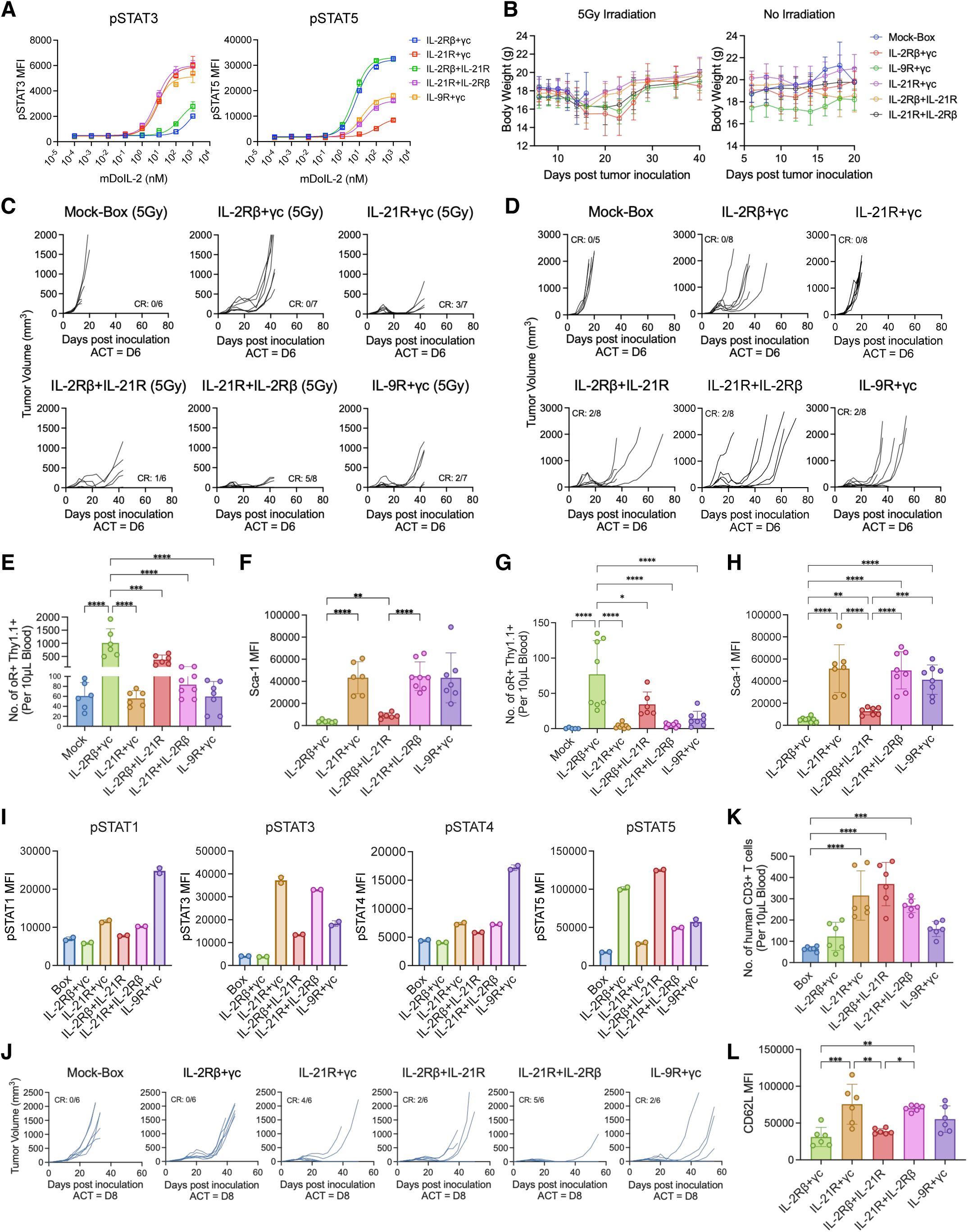
Characterization of enhanced adoptive cell therapy mediated by IL-2Rβ–IL-21R heterodimers in mouse and human tumor models, related to Figure 3. (A) Dose-response curves of pSTAT3 and pSTAT5 signaling for the indicated fully murine orthogonal receptor pairings with mouse DoIL-2 stimulation. (B) Relative body weight change in mice following the indicated ACT treatments. See Figures 3A and 3D for experimental conditions. (C-D) Individual B16F10 tumor growth curves of C57BL/6 mice receiving the indicated ACT treatment, with (C) or without (D) irradiation. Numbers of complete responses (CR) relative to the total number of mice per group are indicated. (E-F) Transferred T cell phenotyping in lymphodepleted syngeneic ACT model. (E) Transferred Pmel-1 T cell counts in 10 μL tail blood at day 7 post-ACT. (F) Sca-1 expression on transferred T cells after DoIL-2 dosing. (G-H) Transferred T cell phenotyping in non-lymphodepleted syngeneic ACT model. (G) Transferred Pmel-1 T cell counts in 10 μL tail blood at day 7 post-ACT. (H) Sca-1 expression on transferred T cells after DoIL-2 dosing. (I) Bar plots showing pSTAT1/3/4/5 activation in transduced primary human T cells expressing the indicated orthogonal receptor pairings following stimulation with 100 nM human DoIL-2. Data are presented as median fluorescence intensity (MFI). (J) Individual A375 tumor growth curves of NSG mice receiving ACT of the indicated human CD3⁺ T cells. Numbers of complete responses (CR) relative to the total number of mice per group are indicated. (K-L) Transferred T cell phenotyping in Human xenograft model. (K) Human CD3+ T cell counts in 10 μL tail blood at day 7 post-ACT. (L) CD62L expression on transferred T cells.

**Figure S4.**
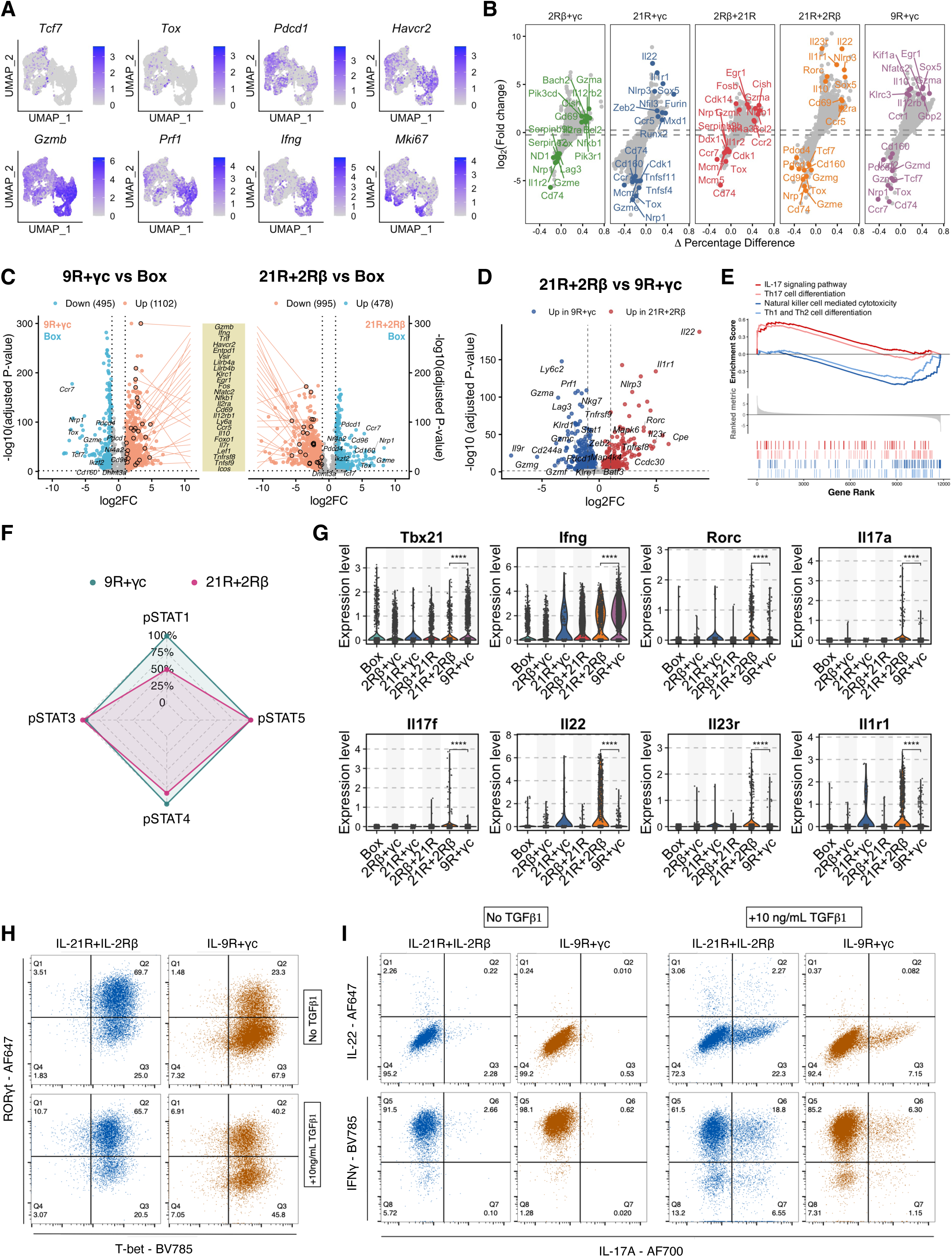
scRNA-seq analysis and phenotypic characterization of tumor-infiltrating Pmel-1 T cells programmed by IL-21R/IL-2Rβ and IL-9R/γc pairings, related to Figure 4. (A) UMAP visualization of gene expression for selected markers associated with T cell functional states. Expression levels of *Tcf7*, *Tox*, *Pdcd1*, *Havcr2*, *Gzmb*, *Prf1*, *Ifng*, and *Mki67* are shown. Color intensity indicates normalized expression levels. (B) Scatter plots showing differential gene expression across the indicated orthogonal receptor pairings relative to Box control. The x-axis represents the difference in the percentage of expressing cells (Δ percentage), and the y-axis represents log₂ fold change (log₂FC). Each point represents a gene, with selected genes labeled. Dashed lines indicate thresholds for differential expression (|log₂FC| > 0.5). (C) Comparative volcano plots showing differentially expressed genes for IL-21R/IL-2Rβ and IL-9R/γc, each versus Box control. Shared genes are connected by lines. (D) Volcano plot of direct IL-21R/IL-2Rβ versus IL-9R/γc comparison, highlighting genes selectively upregulated in each condition. (E) GSEA plots for IL-17 signaling, Th17 cell differentiation, NK cell–mediated cytotoxicity, and Th1/Th2 cell differentiation, based on ranked differential expression between the two conditions. (F) Radar plot comparing relative STAT signaling activation between IL-21R/IL-2Rβ and IL-9R/γc heterodimers. Activation levels of pSTAT1, pSTAT3, pSTAT4, and pSTAT5 are shown. Values are normalized to the maximum response. (G) Violin plot showing the expression levels of Tc1- and Tc17-associated genes across the indicated orthogonal receptor pairings from the scRNA-seq data. Expression of *Tbx21* and *Ifng* (Tc1), and *Rorc*, *Il17a*, *Il17f*, *Il22*, *Il23r*, and *Il1r1* (Tc17) are shown. Statistical significance was determined by one-way ANOVA with multiple comparisons. (H-I) Representative flow cytometry plots with quadrant gating showing intracellular staining of Tc1- and Tc17-associated transcription factors (H) and cytokines (I) under different stimulation conditions, related to Figure 4I and 4J.

**Figure S5.**
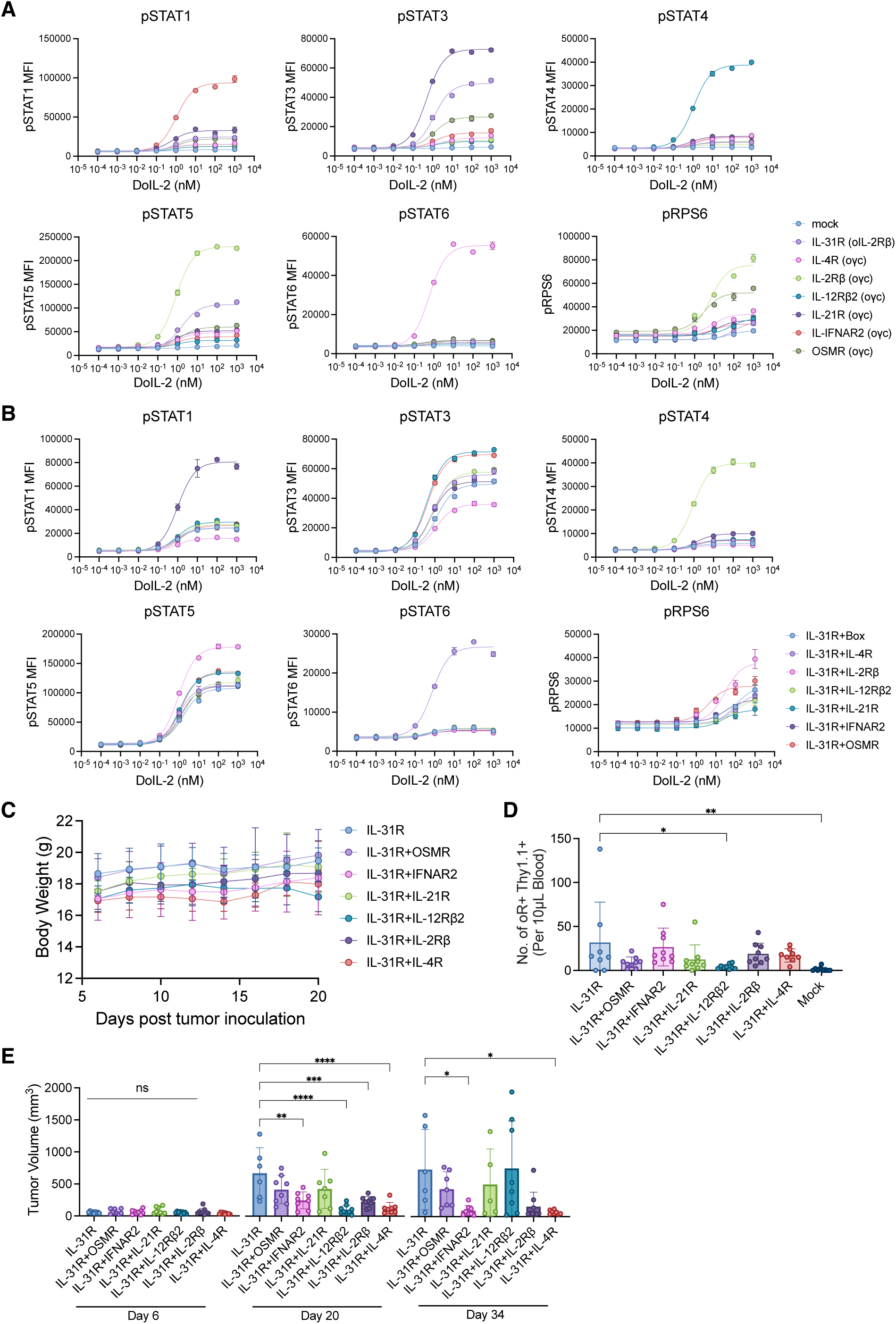
Signaling characterization of IL-31R-based orthogonal receptor pairings, related to Figure 5. (A) Dose-response curves of STAT and RPS6 phosphorylation induced by indicated orthogonal receptor pairings containing a single intracellular domain on one chain, related to Figure 5B. (B) Dose-response curves of STAT and RPS6 phosphorylation induced by the indicated orthogonal receptor pairings containing intracellular domains on both chains, related to Figure 5C. (A-B) Pmel-1 T cells were stimulated with DoIL-2 for 30 min. All data represent mean ± SD. (C) Relative body weight change in mice following the indicated ACT treatments. See Figures 5I for experimental conditions. (D) Number of transduced Pmel-1 CD8⁺ T cells in 10 μL of tail blood 7 days after ACT. See Figures 5I for experimental conditions. Statistical significance was determined by one-way ANOVA followed by Tukey’s multiple comparisons test. *p < 0.05; **p < 0.01. (E) Tumor burden assessed at baseline and at 2 and 4 weeks post-ACT.

**Figure S6.**
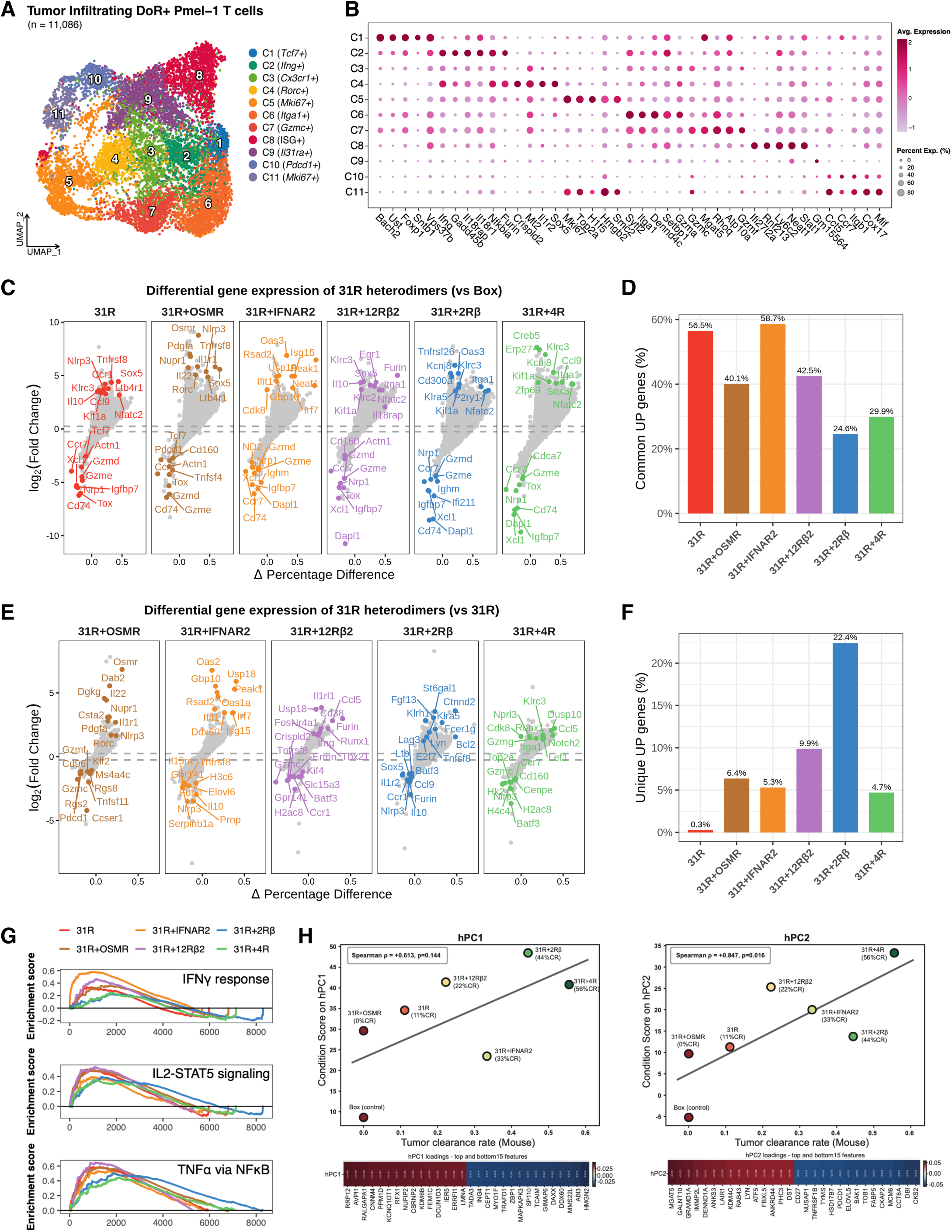
Comparative analysis of the transcriptional landscape of TILs reprogrammed by IL-31R-based pairings, related to Figure 6. (A) UMAP of 11,086 Pmel-1 TILs from B16F10 tumors across IL-31R-based pairing conditions and Box control. Eleven clusters are color-coded with marker gene annotations. (B) Dotplot showing the top 5 marker genes for each cluster identified by differential expression analysis. Dot size represents the percentage of cells expressing each gene, and color intensity indicates normalized expression levels. (C) Scatter plots showing differential gene expression across the indicated IL-31R-based orthogonal receptor pairings relative to Box control. The x-axis represents the difference in the percentage of expressing cells (Δ percentage), and the y-axis represents log₂ fold change (log₂FC). Each point represents a gene, with selected genes labeled. Dashed lines indicate thresholds for differential expression (|log₂FC| > 0.5). (D) Bar plot showing the proportion of the 599 common upregulated genes among the total differentially expressed genes (DEGs) for each indicated sample relative to Box control. (E) Scatter plots showing differential gene expression across the indicated IL-31R-based orthogonal receptor pairings relative to IL-31R alone. The x-axis represents the difference in the percentage of expressing cells (Δ percentage), and the y-axis represents log₂ fold change (log₂FC). Each point represents a gene, with selected genes labeled. Dashed lines indicate thresholds for differential expression (|log₂FC| > 0.5). (F) Bar plot showing the proportion of sample-specific differentially expressed genes (DEGs) among the total DEGs for each indicated sample relative to Box control. (G) Overlaid GSEA enrichment curves for IFN-γ response, IL-2-STAT5 signaling, and TNFα signaling via NF-κB, shown for all six IL-31R-based pairings versus Box control. (H) Correlation between tumor clearance rates and projection scores along hPC1 (left) and hPC2 (right) for IL-31R-based orthogonal receptor pairings and Box control. Spearman correlation coefficients are indicated. The top and bottom 15 human feature genes contributing to each principal component are shown below.

**Figure S7.**
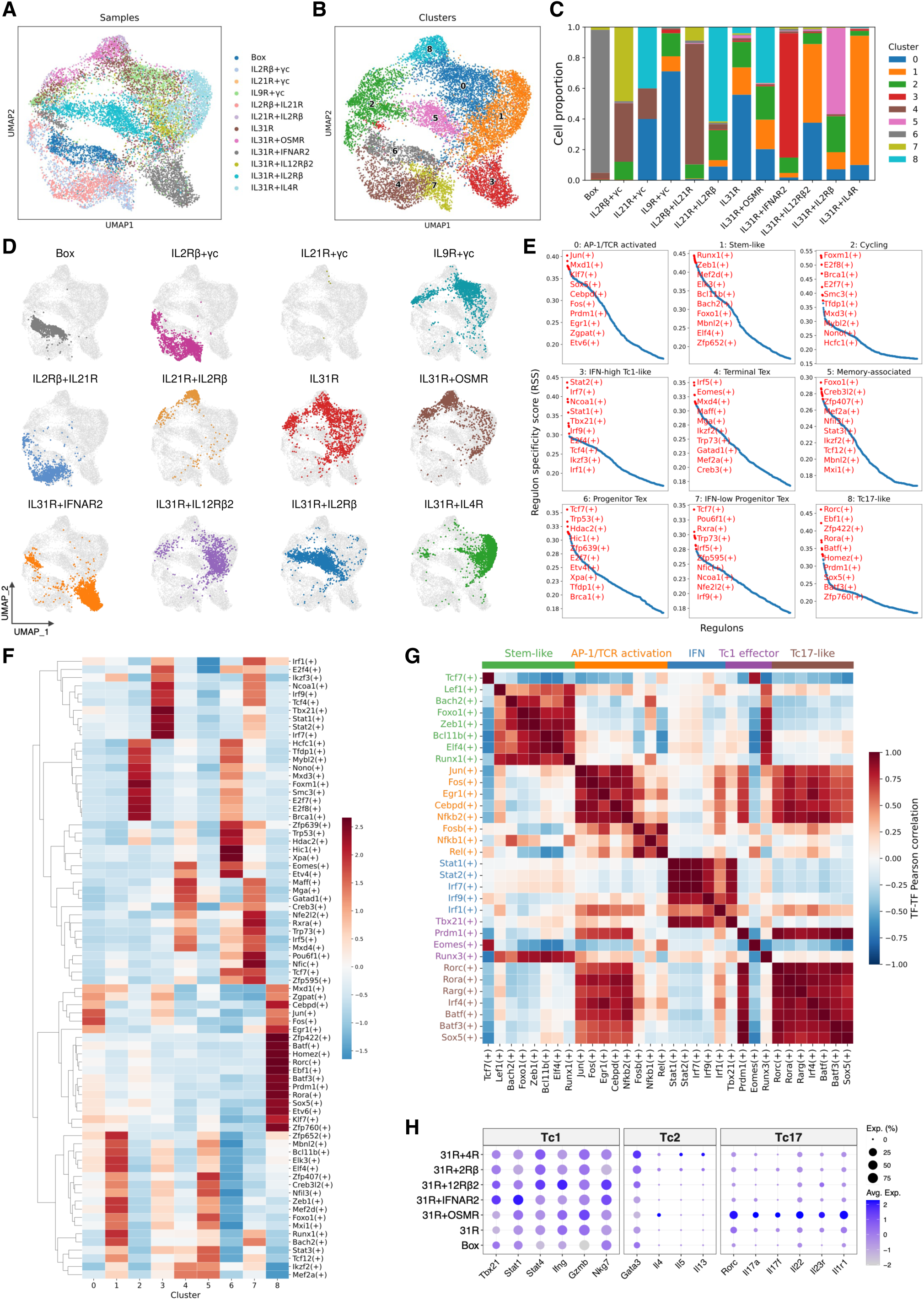
Integrated scRNA-seq and regulon analysis reveal distinct T cell states driven by rewired cytokine receptor pairings, related to Figure 7. (A) UMAP visualization showing sample-level cell distribution in the integrated scRNA-seq dataset. Twelve samples are color-coded, with annotations shown in the right panel. (B) UMAP visualization showing cluster-level cell distribution in the integrated scRNA-seq dataset. Nine major clusters are defined and annotated on the UMAP. (C) Stacked bar plot showing the proportion of cells in each annotated cluster from (B) for each indicated sample, colored by cluster identity. (D) Per-condition UMAPs showing cell distributions for each pairing. (E) Regulon specificity score (RSS) ranking plots showing cluster-specific regulons. Top 10 regulons per cluster are highlighted in red, and clusters are annotated based on top marker regulons representing major T cell functional states. (F) Heatmap showing regulon activity across clusters. Top ten regulons were selected based on regulon specificity scores (RSS) for each cluster. Data represent normalized regulon activity (Z-score). (G) Heatmap showing pairwise correlations of regulon activities across all tested pairings, with regulons grouped into functional states as defined in Figure 7B. (H) Dot plot showing the activity of Tc1-, Tc2-, and Tc17-associated gene signatures across the indicated IL-31R-based receptor pairings. Dot size represents the percentage of cells expressing each signature, and color intensity indicates normalized activity levels.

